# The post-acute phase of SARS-CoV-2 infection in two macaques species is associated with signs of ongoing virus replication and pathology in pulmonary and extrapulmonary tissues

**DOI:** 10.1101/2020.11.05.369413

**Authors:** Kinga P. Böszörményi, Marieke A. Stammes, Zahra C. Fagrouch, Gwendoline Kiemenyi-Kayere, Henk Niphuis, Daniella Mortier, Nikki van Driel, Ivonne Nieuwenhuis, Richard A. W. Vervenne, Tom Haaksma, Boudewijn Ouwerling, Deborah Adema, Roja Fidel Acar, Ella Zuiderwijk-Sick, Lisette Meijer, Petra Mooij, Ed J. Remarque, Herman Oostermeijer, Gerrit Koopman, Alexis C. R. Hoste, Patricia Sastre, Bart L. Haagmans, Ronald E. Bontrop, Jan A.M. Langermans, Willy M. Bogers, Ivanela Kondova, Ernst J. Verschoor, Babs E. Verstrepen

## Abstract

The post-acute phase of SARS-CoV-2 infection was investigated in rhesus macaques (*Macaca mulatta*) and cynomolgus macaques (*Macaca fascicularis*). During the acute phase of infection, SARS-CoV-2 was shed via nose and throat, and viral RNA was occasionally detected in feces. This phase coincided with a transient change in systemic immune activation. Even after the alleged resolution of the infection, as suggested by the absence of viral RNA in nasal and tracheal swabs, computed tomography (CT) and positron emission tomography (PET)-CT were able to reveal pulmonary lesions and activated tracheobronchial lymph nodes in all animals. Post-mortem histological examination of the lung tissue revealed mostly marginal or resolving minimal lesions that were indicative of SARS-CoV-2 infection. Evidence for SARS-CoV-2-induced histopathology was also found in extrapulmonary tissue samples, like conjunctiva, cervical and mesenteric lymph nodes.

However, 5-6 weeks after SARS-CoV-2 exposure, upon necropsy, viral RNA was still detectable in a wide range of tissue samples in 50% of the macaques and included amongst others the heart, the respiratory tract and surrounding lymph nodes, salivary gland, and conjunctiva. Subgenomic messenger RNA was detected in the lungs and tracheobronchial lymph nodes, indicative of ongoing virus replication during the post-acute phase. These results could be relevant for understanding the long-term consequences of COVID-19 in humans.

**Author summary:** More than a year after the start of the pandemic, the long-term consequences of SARS-CoV-2 infection start to surface. The variety of clinical manifestations associated with post-acute COVID-19 suggests the involvement of multiple biological mechanisms. In this study, we show that rhesus and cynomolgus macaques shed virus from their respiratory tract, generate virus-specific humoral immune responses, and show signs of SARS-CoV-2-induced lung pathology. PET-CT revealed that both species showed ongoing mild to moderate pulmonary disease, even after the virus was no longer detectable in nasal and tracheal swabs. Five to six weeks after infection, necropsy confirmed minimal to mild histopathological manifestations in various tissues, like the lungs, heart, lymph nodes, and conjunctiva. We detected Viral RNA in the heart, respiratory tract, and tracheobronchial lymph nodes, and subgenomic messenger RNA in the lungs and surrounding lymph nodes, indicative of ongoing virus replication. We show widespread tissue dissemination of SARS-CoV-2 in infected macaques and the presence of replicating virus in lungs and surrounding lymph nodes after alleged convalescence of infection. This finding is intriguing in the light of long-COVID disease symptoms seen in humans as it has been hypothesized that persistent infection may contribute to this phenomenon.

## Introduction

Biomedical and clinical researchers have tried to define the different phases of a severe acute respiratory syndrome coronavirus 2 (SARS-CoV-2) infection in humans from various perspectives [1-4]. The general consensus is that the initial phase of infection is viremia or the progressive state in which immune responses determine if the person remains healthy or not. This phase, which takes about a week, is followed by the pneumonia phase that generally lasts no longer than three weeks after the initial infection. Then, a convalescent phase is initiated during which the infection resolves, and patients recover from the disease. Nevertheless, evidence is accumulating for the existence of an enlengthened recovery process, even for non-hospitalized patients [5-7]. Moreover, in this phase, chronic symptoms and possible recurrence of disease can be detected, known as long-COVID or post-acute COVID [7, 8]. However, it is still unknown how long this recovery phase lasts.

To halt and control the coronavirus disease 2019 (COVID-19) pandemic, enormous efforts have been initiated worldwide to develop vaccines and antiviral compounds. To achieve these challenging goals, as well as to acquire fundamental knowledge of understanding the mode of infection and its associated pathologies, animal studies play a crucial role.

Various animal species have already proven their value in SARS-CoV-2 research, like mice, hamsters, and ferrets [9-17]. But notwithstanding the great importance of rodent and ferret models, especially non-human primates (NHPs) have shown to play a pivotal role in COVID-19 research. NHPs are susceptible to infection with SARS-CoV-2 and share many immunological and pathological characteristics with humans [9, 18, 19]. This makes NHPs particularly suitable for preclinical evaluation of vaccines and antiviral or immunomodulatory compounds against SARS-CoV-2. The two most widely used NHP species in biomedical research in general, and also in COVID-19 research, are rhesus macaques and cynomolgus macaques. Both species had already featured a prominent role in research on the related coronaviruses that caused the SARS and MERS epidemics [20, 21], and are also considered relevant NHP models for preclinical COVID-19 studies [9].

The choice for a specific macaque (sub)species can be crucial [22-26]. Models for AIDS, TB, and influenza illustrated a differential course of infection, as well as disease development, in different macaque species. When a new and complex infectious agent like SARS-CoV-2 emerges, the choice of an NHP species for research is therefore not a trivial one. Cynomolgus macaques have been deployed in studies describing aspects of SARS-CoV-2 pathogenesis [27-29] and have been utilized to evaluate the efficacy of hydroxychloroquine as an antiviral compound [30]. Rhesus macaques have also been applied in COVID-19 pathogenesis studies [31-34], and to test antiviral compounds [35]. Many COVID-19 vaccines, among others those already authorized or approved for use in humans, have received their first efficacy evaluation in the rhesus macaque model [12, 36-45].

Several research groups [27, 32, 46] shed light on the heterogeneity in SARS-CoV-2 infection and investigated disease progression in different NHP species, but a controlled comparative approach is lacking. The key question is which macaque species is best suited for the investigation of certain aspects of SARS-CoV-2 and COVID-19. To address this issue, we directly compared SARS-CoV-2 replication in rhesus and cynomolgus macaques and monitored the animals for signs of COVID-19-like disease symptoms. The macaques were infected in parallel with the same virus stock and the same dose and underwent a fully identical treatment. The course of infection was followed for up to six weeks using the same analyses, including monitoring of virus-induced metabolic and anatomic findings with CT and PET-CT, and continuous telemetric recording of body temperature and physical activity of the animals. The animals were sacrificed to investigate histopathological tissue changes related to SARS-CoV-2, as well as to examine viral dissemination in organs.

Until recently, only short follow-up studies were published, sacrificing animals at day 21 post-infection, or earlier [27, 29, 47, 48]. Here, we took a unique approach and focused our research on the first weeks of the post-acute phase, after a mild to moderate SARS-CoV-2 disease course in two macaque species.

## Results

### Infection of macaques with SARS-CoV-2

Following the administration of the virus in the upper trachea and nose, viral RNA was detectable in the tracheal and nasal swabs of all monkeys at day 1 post-infection (pi). The time frame in which viral RNA was detected varied considerably from one day (tracheal swab macaque R15096) to up to ten days in macaque R14002. (Fig 1A-B and D-E, Table S1A). The individual variation of SARS-CoV-2 RNA levels detected in the macaques was, regardless of species, substantial. Peak viral RNA levels in the throat varied between 1.7 x 10^4^ genome equivalent (GE)/ml (R15096; day 1 pi) and 1.8 x 10^8^ GE/ml (J16017; day 2 pi). Peak viral loads detected in nasal swabs were generally lower than in the throat samples and did not exceed 9.5 x 10^4^ GE/ml (R15090; day 1 pi). The high RNA loads measured in the first two days post-infection are suggestive of residual RNA from the original inoculum still being present. However, nasal swabs from five out of eight animals tested positive again after one or more days of undetectable levels. Cynomolgus macaque J16017 was positive in the nose at day 1 pi, then had no detectable viral RNA for a period of three days, but later the animal became again positive in the nose swabs for three consecutive days. Furthermore, R15096, J16004, J16012, and Ji408005 became PCR-positive again after one or more days without detectable viral RNA. The total viral RNA production detected in nose and throat samples was comparable over time for both species (Fig 1G).

**Figure 1.**
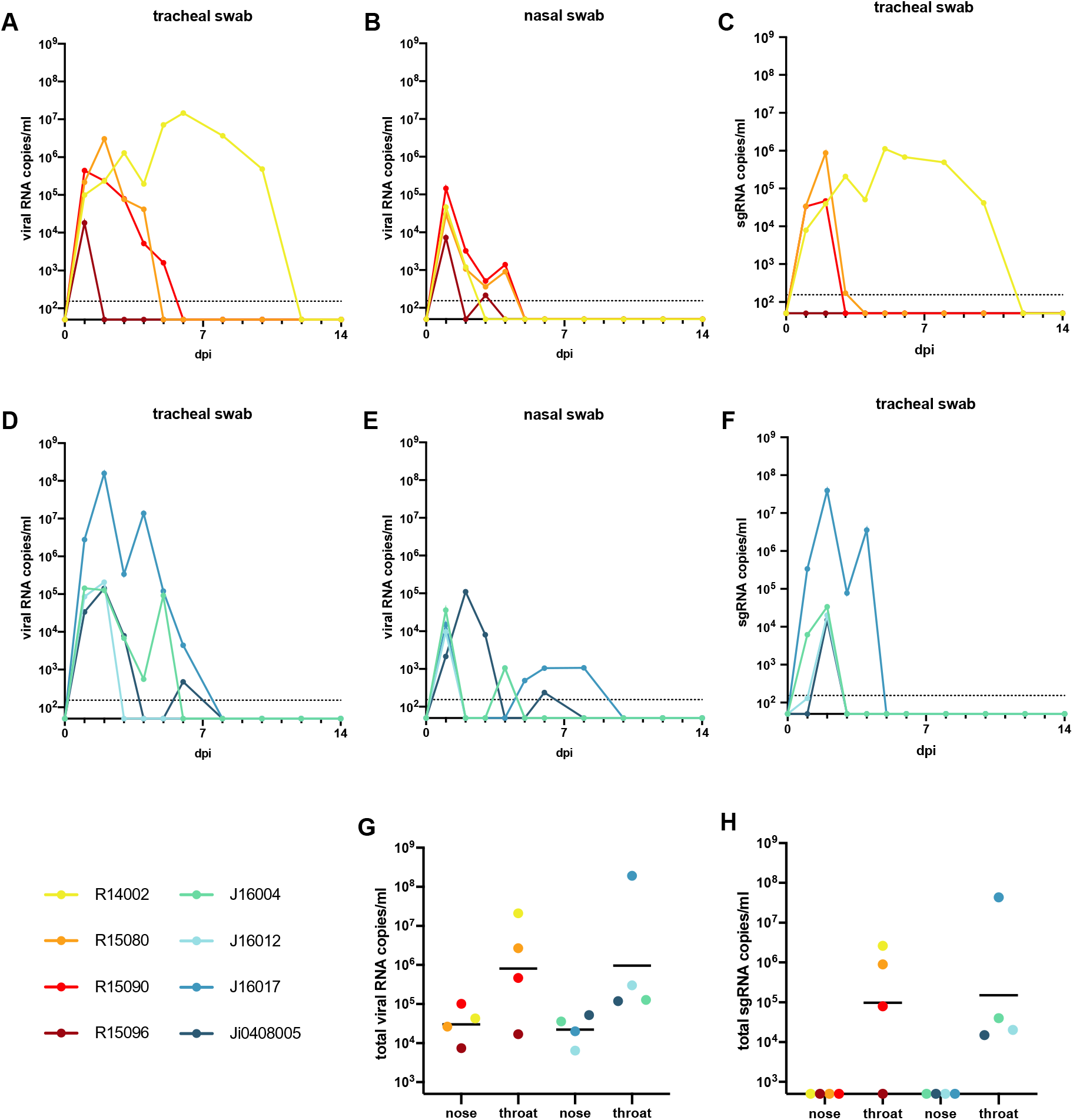
Virus loads in swab samples of macaques following SARS-CoV-2 infection. Viral RNA and subgenomic messenger RNA quantification in tracheal and nasal swabs of rhesus and cynomolgus macaques by qRT-PCR. The limit of quantification is indicated by the dotted horizontal line. (A-C). Viral RNA levels in tracheal (A) and nasal swabs (B), and sgmRNA levels in tracheal swabs (C) of rhesus macaques. (D-F). Viral RNA levels in tracheal (D) and nasal swabs (E), and sgmRNA levels in trachea swabs (C) of cynomolgus macaques. The limit of quantification is indicated by the dotted horizontal line. Total virus loads (G) and sgmRNA loads (H) in throat and nose samples of macaques throughout the study. Horizontal bars represent geometric means. The sum of the viral copies was calculated rather than area under the curve (AUC), as AUC interpolates for time points when virus loads were not determined. The color coding used for each individual animal as shown in the figure is used throughout the manuscript to denote the same individual. Rhesus macaques are indicated by yellow to red colors; cynomolgus macaques by green to blue.

The analysis of viral subgenomic messenger RNA (sgmRNA), which is considered to better reflect actual virus replication, confirmed infection in 7 out of 8 macaques. Only rhesus macaque R15096, which already displayed relatively low levels of viral RNA (Fig 1A-B) was negative for sgmRNA in both throat and nose swabs at all time points. In the remaining animals, sgmRNA levels were exclusively detected in the tracheal swabs, while no evidence of replication was found in the nasal swab samples (Fig 1C, F and H, Table S1B).

In the anal swabs, viral RNA was rarely detected. Few macaques irregularly tested positive in the PCR test, with a low maximum viral RNA load of 3 x 10^3^ GE/ml at day 1 pi (J16017). No sgmRNA was detected in anal swabs or blood samples, indicating the absence of active virus replication in blood and intestinal tract in the animals.

### Body temperature, activity, clinical symptoms and blood parameters after SARS-CoV-2 infection

Body temperature and activity of each animal were continuously monitored using telemetry during the entire study. Modestly elevated body temperatures were measured in both macaque species during the first two weeks after infection as compared to later time points. But whereas three out of four cynomolgus macaques peaked as early as day 2 pi, the elevation in body temperature in two out of four rhesus macaques occurred later on day 8 pi. The temperature curves for the individual animals, depicted in the supplementary figure S1, show a return to baseline temperatures for all individuals after the initial phase of infection.

The cumulative activity scores per week were calculated as the area under the curve (Fig 2). A significantly lower activity during the first week compared to the third week after infection was observed in the rhesus macaques (paired t-test; p=0.0061) suggesting an impact of SARS-CoV-2 infection on the general well-being of the animals. Notably, in the last 2 weeks of the study, activity scores decreased again for all rhesus macaques (p=0.0193). In contrast with these observations, the activity scores of the cynomolgus macaques remained unaffected throughout the study, which implies a species-related difference in reaction to SARS-CoV-2 infection. Activity curves for the individual animals are documented in supplementary figure S2.

**Figure 2.**
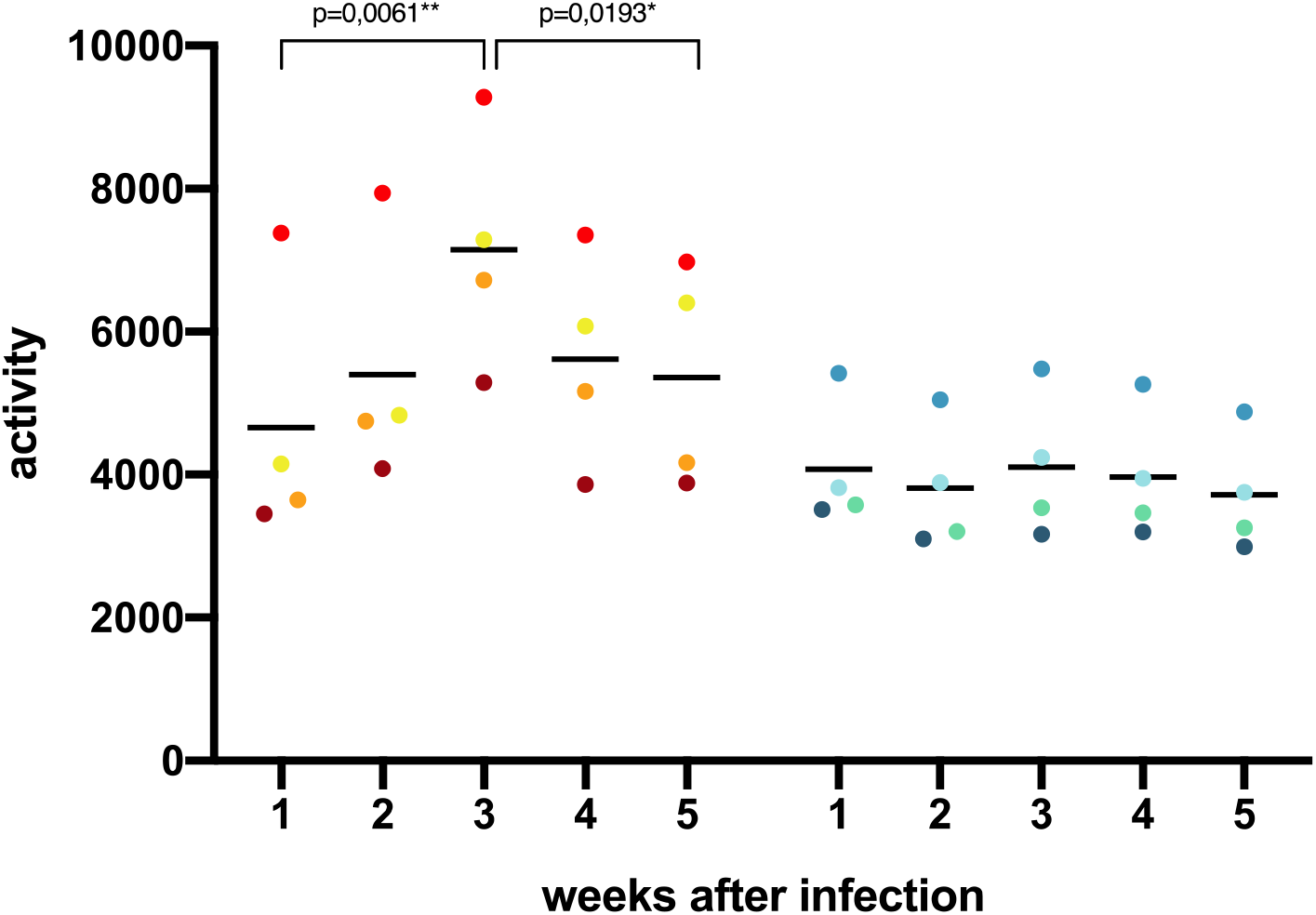
Cumulative activity scores. The activity of the animals was measured by telemetry throughout the study. The cumulative activity scores for rhesus and cynomolgus macaques were calculated area under the curve (AUC) per week. Measurements done during biotechnical handlings were omitted from the calculations.

A scoring list was used to enumerate overt clinical symptoms that may be caused by the SARS-CoV-2 infection (Table S2). The cumulative clinical score per week of each animal was calculated (Fig S3). In the first week after infection, the scoring of the four rhesus and two cynomolgus macaques did not exceed 25 (of maximum 770 per week), while for the remaining two animals a higher, but statistically not significant, median score was calculated. In the subsequent weeks after infection, none of the animals showed manifest clinical signs suggestive of COVID-19.

Blood samples were analyzed for changes in cell subsets and in biochemical parameters upon SARS-CoV-2 infection (Fig S4). Data were related to a set of normal (standard) values derived from a large group of uninfected, healthy macaques from the same breeding colony. C-reactive protein levels, which are increased in COVID-19 patients with pneumonia [49], were not elevated in the infected macaques. In humans, acute kidney injury has been related to SARS-CoV-2 infection [50, 51], and elevated levels of serum creatinine and blood urea were detected in 10-15% of a cohort of COVID-19 patients [52]. Hence, we measured creatinine and urea levels in blood samples at multiple days post-infection but did not find evidence of kidney malfunction in the macaques. Equally, depending on the severity of the disease, blood coagulation disorders, like highly elevated D-dimer levels, have been reported for patients [53, 54], but no elevated D-dimer levels were measured in either macaque species. Elevated levels of glucose and alanine transferase were measured in the first week pi in the blood of most animals, and amylase was increased in one rhesus macaque, R15080. Also, for other blood cell subsets, no significant deviations from the normal values were seen in the infected monkeys.

### Humoral immune response to SARS-CoV-2 infection

Humoral immune responses to the viral spike (S) and nucleoprotein (N) proteins were readily detectable after infection (Figs 3 and S5).

IgG titers to the ectodomain of the spike protein were first detectable 6 to 8 days pi. Titers continued to rise and reached a plateau after approximately 4 weeks. IgG levels in cynomolgus macaques showed more variation compared to rhesus macaques. Especially, the IgG kinetics in J16012 deviated from the other animals. In this animal, a peak in IgG was observed 17 days pi, and titers rapidly declined thereafter, and then remained consistently low throughout the rest of the study period. Overall, IgG was detectable slightly earlier in cynomolgus macaques compared to rhesus macaques, but the differences were not statistically relevant. In addition to the total anti-S IgG, IgG directed to the receptor-binding domain (RBD) were analyzed. The development of anti-RBD IgG fully mirrored the IgG antibodies binding to the entire S protein. IgM directed to the full-length S protein and the RBD were also measured (Fig S5), and titers developed around the same time point post-infection as IgG titers. Overall, IgM titers in cynomolgus macaques were higher than in rhesus macaques, except for R15090. Anti-RBD IgM in rhesus macaques remained low in the study period, while in 3 out of 4 cynomolgus macaques rising anti-RBD IgM titers were measured.

In addition to the immunoglobulin responses to the spike protein, antibody responses to the nucleoprotein (N) were measured. Total immunoglobulin (Ig) in sera was determined by DR-ELISA. Ig responses became evident between day 10 and 12 pi. For both rhesus as well as cynomolgus macaques, a peak in levels was measured between 12 and 23 days pi, except in Ji0408005, where the peak was at day 30 pi.

Total Ig was reflected by IgG directed to the N protein. Notably, IgM responses to N were virtually undetectable in the longitudinal serum samples. Only in one animal, cynomolgus macaque J16012, IgM titers were transiently detectable (Fig S5).

### Cytokine and chemokine measurements in sera of infected macaques

We measured a panel of 13 cytokines and chemokines to characterize the inflammatory response triggered by SARS-CoV-2. In general, the cytokine and chemokine responses reflect two phases. The first phase is characterized by a transient decrease in IP-10, IL-6, MIP-1α, MIP-1β, and IFN-γ levels (Fig 4; Fig S6 individual levels) while others were maintained at a stable level (Eotaxin, TNF-*α*, and IL-8). This trend was observed in both species, except for IP-10 and RANTES. At day 2 pi, the serum levels of IP-10 were 9-fold (95% CI 2.72 - 29.89) higher in the cynomolgus macaques than in rhesus macaques (t-test; p = 0.005). Also, the response pattern of RANTES was different between the two species. In cynomolgus macaques, RANTES rapidly decreased after exposure to SARS-CoV-2 and slowly returned to normal levels at the end of the follow-up period. In rhesus macaques, a transient dip in RANTES levels was observed early after infection. The levels returned to normal within 8 days pi but started to decline again in de following days. At the end of the study, RANTES levels were back to normal in both species. MCP-1 (CCL2) showed a variable increase in 2 out of 4 rhesus macaques and in all cynomolgus macaques (Fig S6).

**Figure 3.**
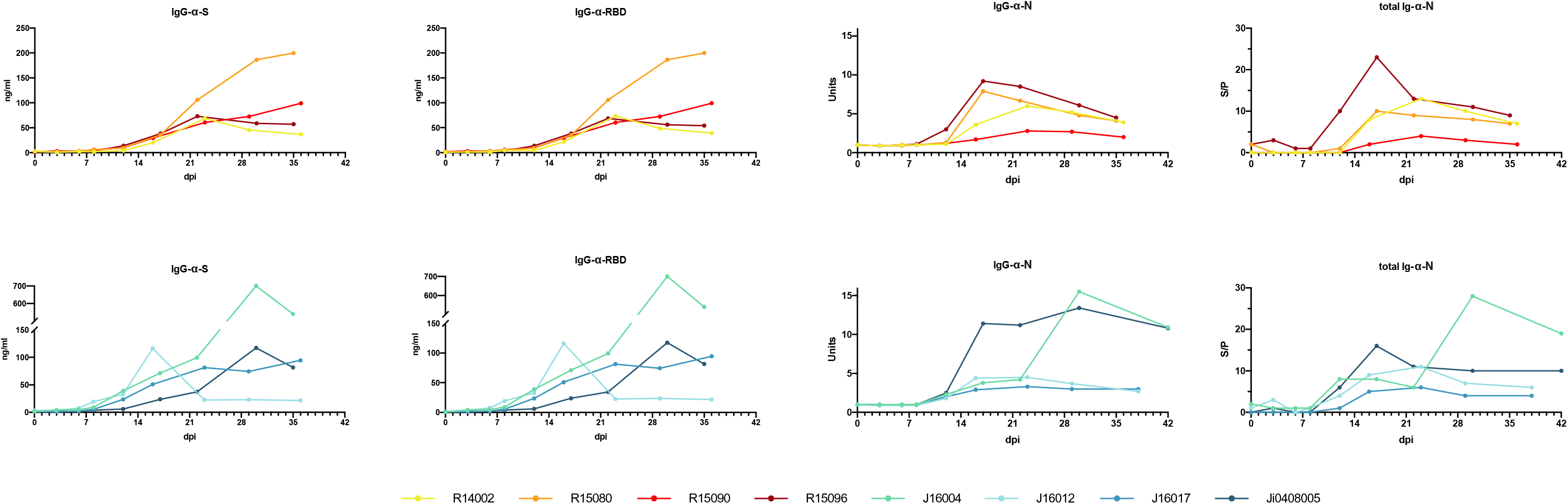
Development of SARS-CoV-2 antibody response in rhesus and cynomolgus macaques. The humoral immune response was determined using an anti-S IgG ELISA, an anti-RBD ELISA, a serological test to detect IgG directed to the N protein, and a DR-ELISA measuring the total antibody response to N (left to right). Results of DR-ELISA are shown as S/P: sample to positive control ratio: 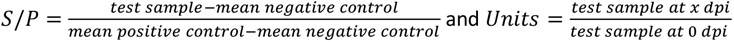

**Figure 4.**
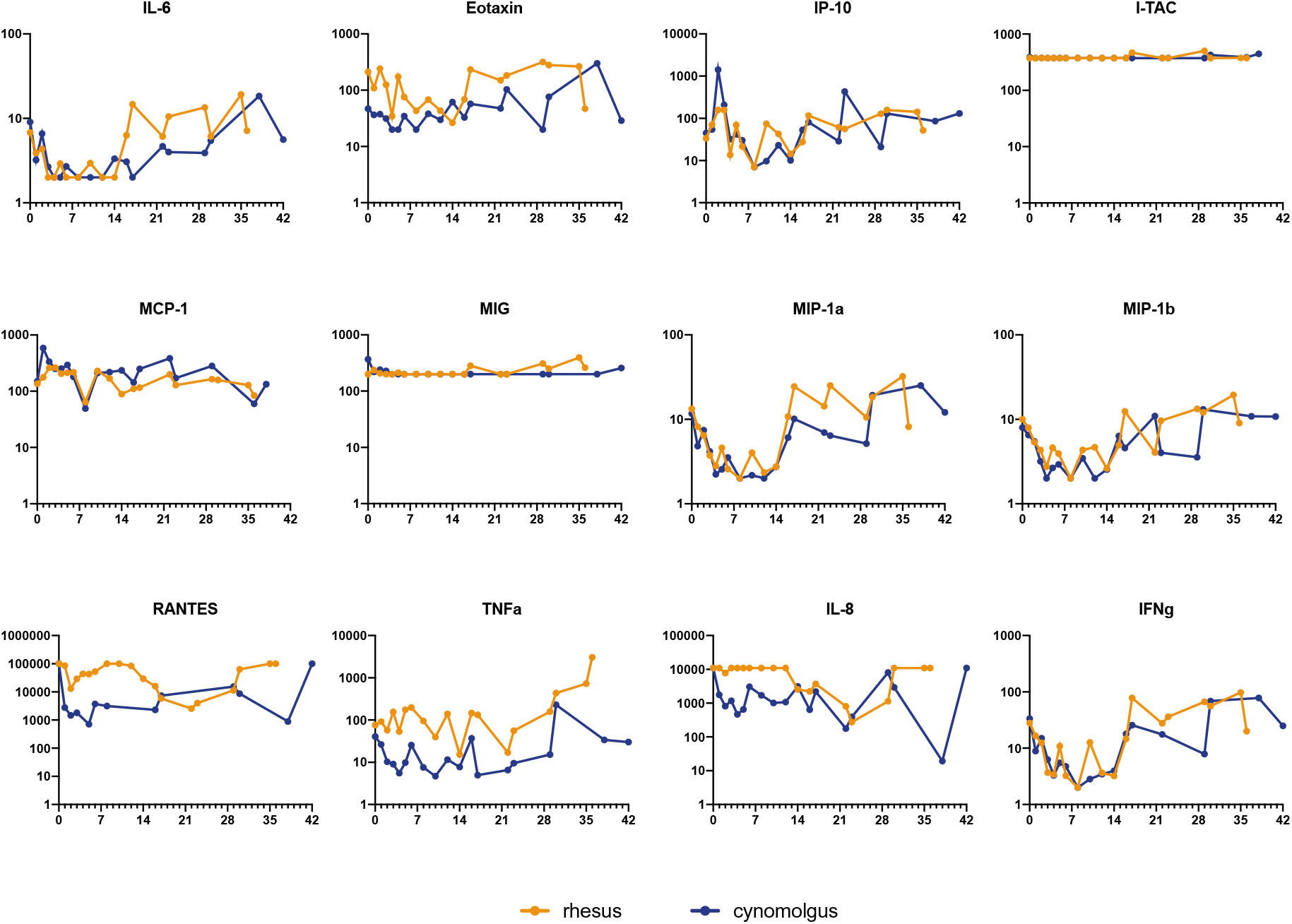
Cytokine and chemokine levels in peripheral blood after SARS-CoV-2 infection. Longitudinal sera collected after SARS-CoV-2 infection were tested. Results were expressed as pg/ml. Statistically significant differences in the geometric mean between rhesus and cynomolgus macaques are indicated with *.

### Development of lung lesions and lymph node activation during SARS-CoV2 infection

Chest CTs of the macaques after infection revealed several manifestations of COVID-19 with a variable time course and lung involvement. The most common lesions found were ground-glass opacities (GGO), consolidations, and crazy paving patterns [3, 55, 56]. Lung lesions were already seen in the first CT obtained two days after infection (max. CT score 2.5/30), in 5 out of 8 monkeys, three rhesus, and two cynomolgus macaques. Thereafter, lung involvement was seen in most animals and CT scores increased. Around days 8 and 10 pi, lung lesions had become manifest in all animals, and in several macaques, the scores had increased (Table 1). Individual differences were considerable, varying from low scores on irregular time points (R15090) to high CT scores almost throughout the entire study period (J16012). The average cumulative CT score at day 38 pi was 12.9 for the rhesus macaques and 25 for the cynomolgus macaques (Mann-Whitney-U test, p=0.1702) with an average increase of 0.44 and 0.71 per day for rhesus and cynomolgus macaques, respectively (Mann-Whitney-U test, p= 0.5036) (Fig 5).

**Figure 5.**
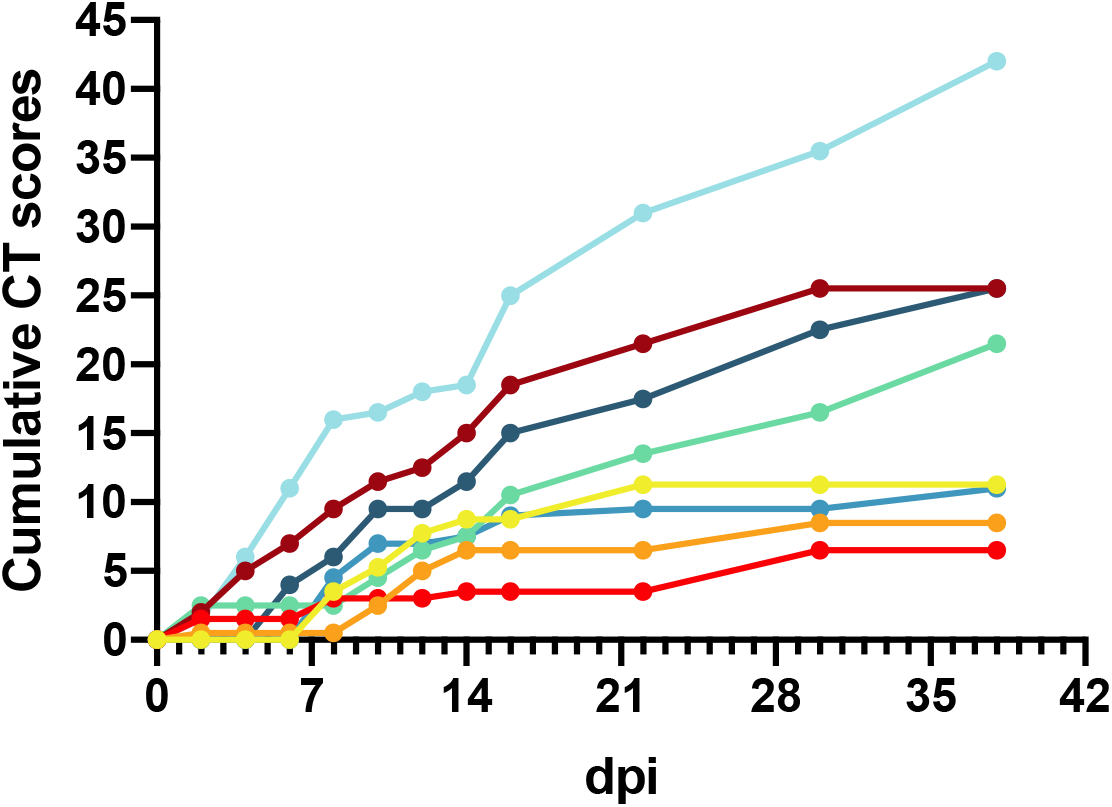
Cumulative CT scores. Cumulative CT scores for each animal were calculated based on the CT scores depicted in Table 1.

**Table 1.** CT scores of lung lesions in SARS-CoV-2-infected macaques. Scoring was performed independently by two experienced scientists.

|  | Animals |  |  |  |  |  |  |  |
| --- | --- | --- | --- | --- | --- | --- | --- | --- |
|  | R14002 | R15080 | R15090 | R15096 | J16004 | J16012 | J16017 | Ji408005 |
| 0 | 0 | 0 | 0 | 0 | 0 | 0 | 0 | 0 |
| 2 | 0 | 0.5 | 1.5 | 2 | 2.5 | 1.5 | 0 | 0 |
| 4 | 0 | 0 | 0 | 3 | 0 | 4.5 | 0 | 0 |
| 6 | 0 | 0 | 0 | 2 | 0 | 5 | 0 | 4 |
| 8 | 3.5 | 0 | 1.5 | 2.5 | 0 | 5 | 4.5 | 2 |
| 10 | 1.75 | 2 | 0 | 2 | 2 | 0.5 | 2.5 | 3.5 |
| 12 | 2.5 | 2.5 | 0 | 1 | 2 | 1.5 | 0 | 0 |
| 14 | 1 | 1.5 | 0.5 | 2.5 | 1 | 0.5 | 0.5 | 2 |
| 16 | 0 | 0 | 0 | 3.5 | 3 | 6.5 | 1.5 | 3.5 |
| 22 | 2.5 | 0 | 0 | 3 | 3 | 6 | 0.5 | 2.5 |
| 29/30 | 0 | 2 | 3 | 4 | 3 | 4.5 | 0 | 5 |
| 36/38 | nd | nd | nd | nd | 5 | 6.5 | 1.5 | 3 |

For visualization and quantification of lymph node activation in the mediastinum, the tracheobronchial area, 4 (rhesus macaques) or 5 (cynomolgus macaques) PET-CTs were obtained over a period of 5 weeks starting at day 8 pi. The PET-CTs showed increased metabolic uptake in at least one tracheobronchial lymph node for all the macaques on all time points. The volume of the lymph nodes associated with this increased uptake ranged from 27 mm^3^ to 2350 mm^3^ (mean 620 mm^3^, std 534 mm^3^) (Fig S7). On average, a lymph node is defined as enlarged when the volume is at least 194 mm^3^ [57]. The volume, average uptake (SUVmean), and the maximum uptake (SUVpeak) of the lymph nodes decreased simultaneously, while the density stayed approximately the same over time for most animals, except for one animal (J16012). This indicates that even in the clearly enlarged lymph nodes no form of necrosis was initiated over time, as otherwise the density would have changed, the maximum uptake would be unaffected, and the average uptake would have decreased. No differences were observed between rhesus and cynomolgus macaques apart from the anatomical density represented by the Hounsfield units (HUs) (p<0.001) (Fig S7). Based on control scans in healthy animals (n=24) we determined a negative correlation (r=0.7154, p=<0.0001) between weight and anatomical density of, at least, the subcarinal area in macaques, which can explain this difference.

From the longitudinal analysis, we conclude that the differentiation of lung lesions can be performed solely on the CT results, as the metabolic activity was only minimally increased on the PET images. An interesting finding is that the location of the lung lesions was not fixed but varied per time point, for instance, in macaque J16012. This animal showed the most prominent lesions in both lungs. The longitudinal PET-CT follow-up (Fig 6) showed two “waves” of the lymph nodes’ activation. The first peak on day 8 and another peak on day 29 of which the explanation is unknown. At the time point of euthanasia (day 35), activation of tracheobronchial lymph nodes was still observed which is indicative for SARS-CoV-2 induced pathogenesis in the post-acute or convalescent phase of the infection.

**Figure 6.**
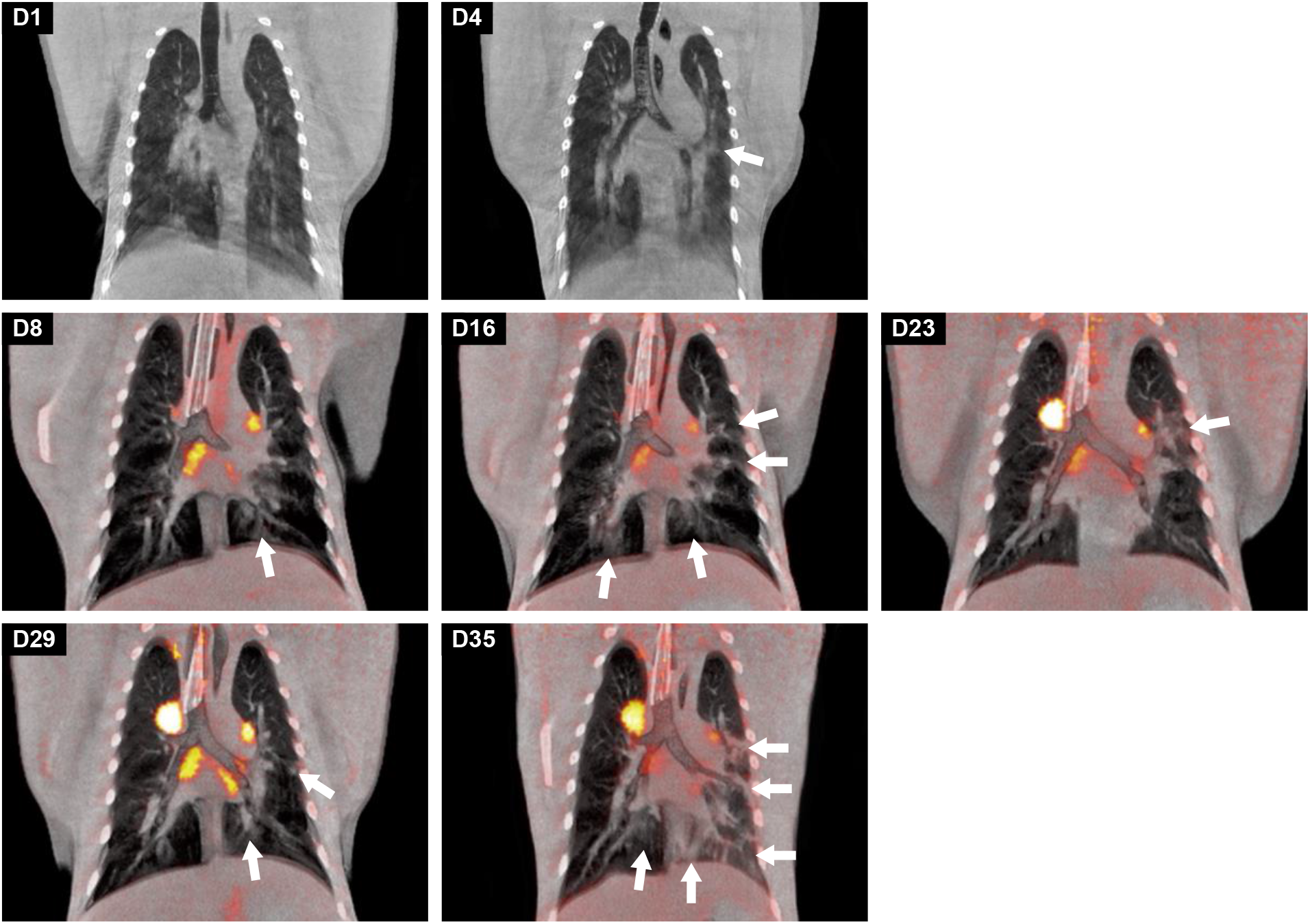
Longitudinal development of both lung lesions and metabolic activity in tracheobronchial lymph nodes of J16017 over time after a SARS-CoV-2 infection. Representative coronal slices with a thickness of 3mm of J16017 were used for visualization. On days 0 and 4, only CT images were obtained, afterwards, until day 35, CT was combined with PET. The location of the lesions, marked with arrows, differed almost per time point, but are most prominently localized in the left lung on day 35.

### Gross pathology

Rhesus macaques R15080 and R15096 were euthanized on day 35 pi, and animals R15090 and R14002, on day 36 pi. The cynomolgus macaques J16012 and J16017 were euthanized on day 38 pi and animals J16004 and Ji0408005 on day 42 pi.

Macroscopically, the lungs of the rhesus and cynomolgus macaques appeared mostly unaffected. One rhesus macaque (R15096) had few small foci with hyperemia at the dorsal aspects of the caudal left and right lung lobes, and in three cynomolgus macaques the lungs had similar foci of hyperemia, confined also to the caudal pulmonary lobes. The gross examination of extrapulmonary organs revealed mildly to moderately enlarged cervical and mesenteric lymph nodes. The rest of the organs were macroscopically unaffected.

### Wide-spread presence of viral RNA in post-mortem tissue samples, and evidence of active virus replication in the respiratory tract of infected macaques

Post-mortem tissue samples were analyzed for the presence of viral RNA (Fig 7, Table S3) and subgenomic messenger RNA (Table 2). After euthanasia, viral RNA was detected in tissue samples of four out of eight animals, one rhesus (R14002) and three cynomolgus macaques (J16012, J16017, and Ji408005). Between the PCR-positive animals striking differences were observed in the number of positive tissues. J16012 and Ji408005 were only positive in respiratory tract tissues, Ji408005 tested positive in the pharyngeal mucosa, lung and two lung lymph nodes, while viral RNA was detected in the carinal part of the trachea, lung and four lung lymph nodes of animal J16012. In the two other PCR-positive macaques (R14002 and J16017), RNA was detected in the respiratory system and the tracheobronchial lymph nodes, but also in cervical and mesenteric lymph nodes, skin, conjunctiva, liver, spleen, kidney, salivary gland and heart, all organs and tissues that have been described as target tissues in humans [58]. High viral RNA loads were measured in some lymph nodes of these animals, ranging from 2 x 10^5^ to 1 x 10^6^ GE/gram of tissue. Also, in the heart of J16017 and R10002, high viral RNA loads (between 5 x 10^3^and 1.15 x 10^5^ GE/gram of tissue) were detected (Table S3).

**Table 2.**
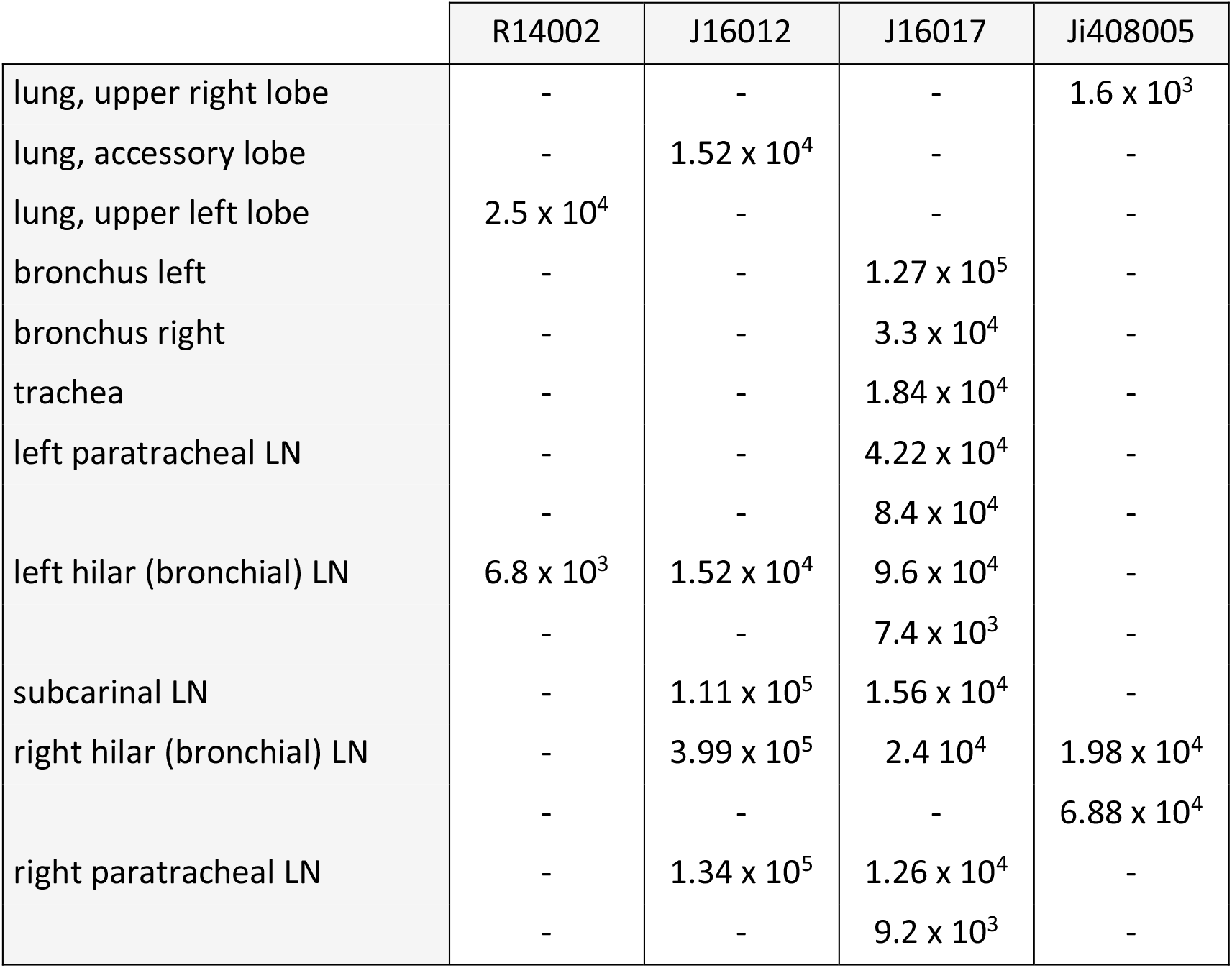
Determination of sgmRNA copies per gram of tissue. PCR analysis was done on all tissues tested positive for SARS-CoV-2 RNA in the RdRp gene assay (Fig 7). Only sgmRNA PCR-positive tissue samples are indicated.

**Figure 7.**
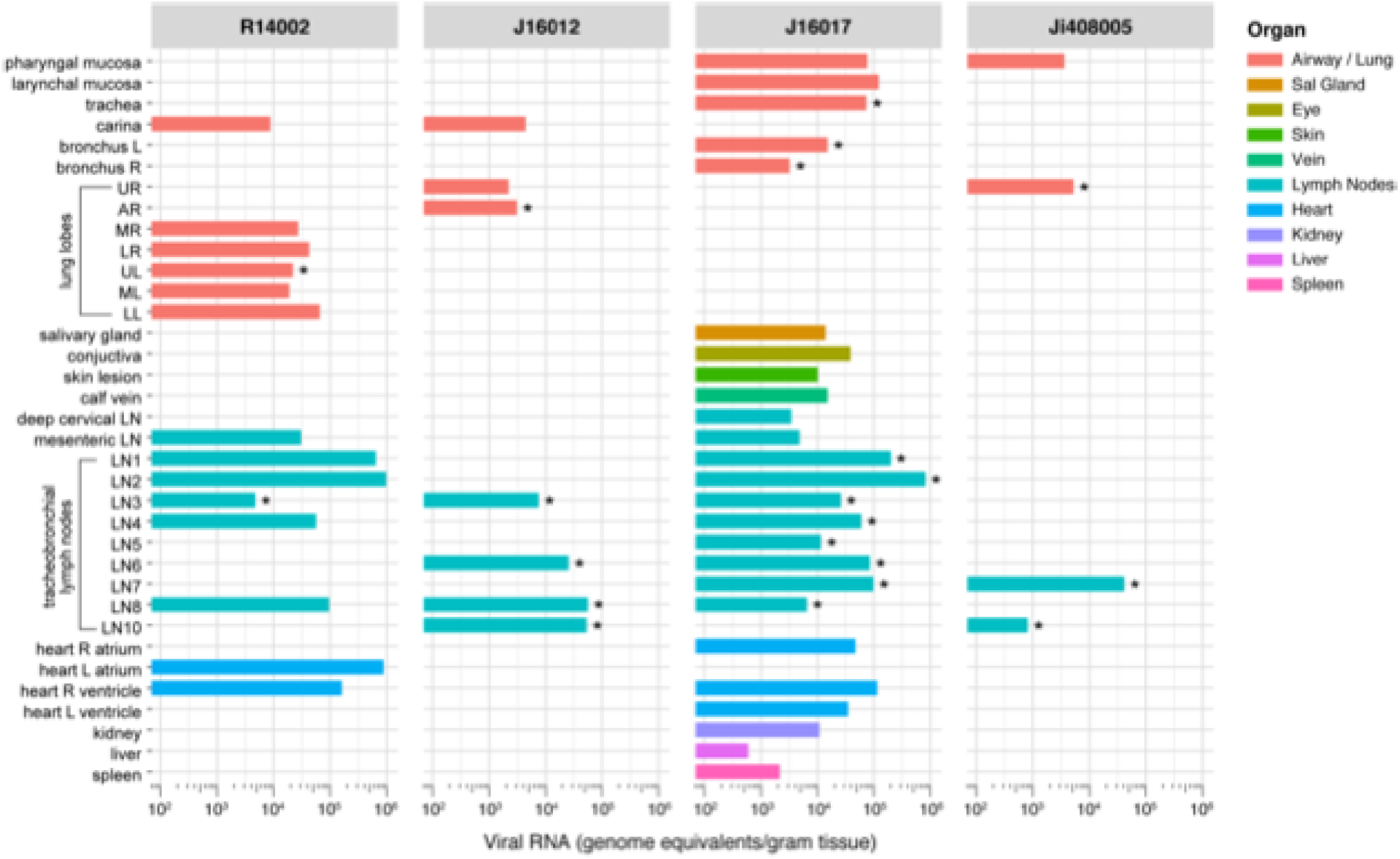
Viral RNA loads in tissues of SARS-CoV-2-infected macaques. Viral RNA in tissues was quantified by qRT-PCR and calculated per gram of tissues. Only animals with viral RNA positive tissues are shown, and only tissues are depicted that were positive in at least one animal. Various (groups of) tissues are indicated by color codes as indicated on the right. Tissues that tested positive for sgmRNA are indicated with *.

Active SARS-CoV-2 replication was evidenced by using sgmRNA PCR analysis on all tissue samples that tested positive in previous viral RNA test. As sgmRNA is synthesized during virus replication, this assay is recognized as an alternative to virus isolation from tissue samples. In table 2 and figure 7, the sgmRNA-positive tissues are shown. Despite the fact that virus was no longer detectable in nasal and tracheal swabs from the 4 animals for more than 26 days (R14002), 30 days (J16017), or even 34 days (J16012 and Ji408005), sgmRNA was detectable in respiratory tract tissues of all four macaques, convincingly indicating that SARS-CoV-2 continued to replicate in these animals after the alleged resolution of infection that was concluded from the negative nasal and tracheal swab samples.

### Histopathology

The histological examination of the lung tissue samples revealed unremarkable or resolving minimal lesions that were indicative of a previous infection with SARS-CoV-2. The nature of the lesions was similar in both rhesus and cynomolgus macaques and comprised of some or all of the following: focal areas with minimal to mild interstitial mononuclear inflammatory infiltrates, foci of bronchiolar smooth muscle hyperplasia (Fig 8A panel 2), vascular congestion, multifocal mild expansion of alveolar septa with delicate collagen (Trichrome stain; Fig 8A panel 3), and occasional foci of type II pneumocytes hyperplasia and few intra-alveolar macrophages (Fig. 8A panel 4). The extrapulmonary organs were histologically mostly unremarkable, or exhibited mild, nonspecific background lesions like minimal interstitial lymphocytic infiltrates, acute-chronic colitis, and enteritis. However, the conjunctiva of cynomolgus macaque J16017 showed multifocal and perivascular foci of moderate lymphocytic infiltrates in the substantia propria (Fig.8C inset). Indeed, this finding was supported by PET-CT images (Fig 8C): an increased FDG-uptake was measured in the left eye compared to the right eye of J16017 that is indicative of ongoing SARS-CoV-2-induced pathogenesis. PCR analysis confirmed the presence of viral RNA in the eye of this animal (Fig 7), clearly demonstrating ocular SARS-CoV-2 infection in J16017. In addition, the cervical lymph node of the same animal exhibited marked lymphoid hyperplasia (follicular type) (Fig 8B panel 1). Similar to the conjunctiva, this histological finding was accompanied by the detection of SARS-CoV-2 RNA in this lymph node.

**Figure 8.**
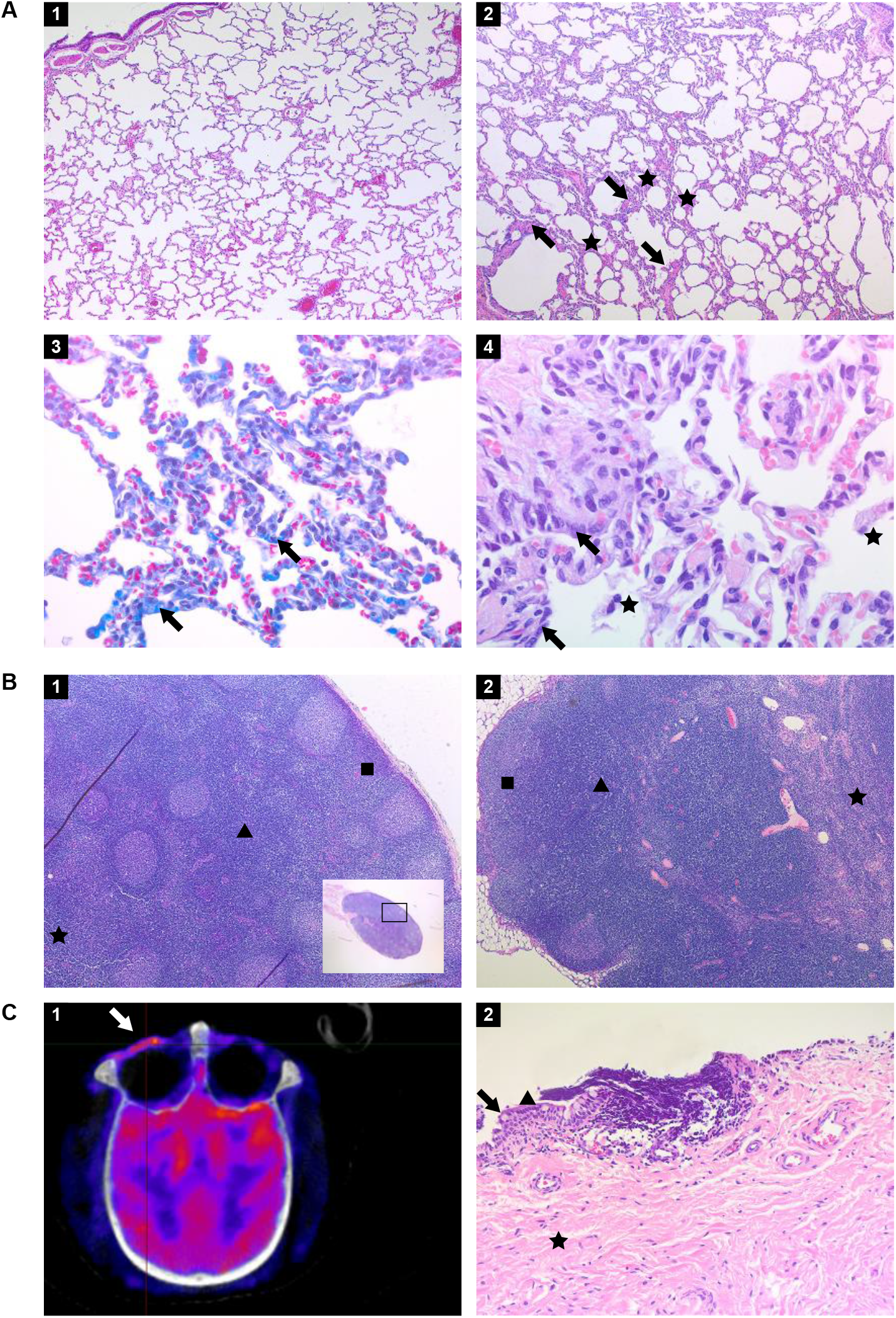
Histopathology of tissues from rhesus and cynomolgus macaques infected with SARS-CoV-2 and euthanized 36 to 42 days post-infection. (A) panel 1. Lung of healthy, uninfected macaque (hematoxylin and eosin (HE) staining, original magnification 50 X); panel 2. Lung of rhesus macaque infected with SARS-CoV-2 and euthanized day 36 pi. Focal area with mild alveolar septa expansion with low number of mononuclear inflammatory cells (asterisks) and foci of bronchiolar smooth muscle hyperplasia (arrows) (HE, 50 X); panel 3. Trichrome staining revealing the presence of collagen mildly expanding the alveolar septa at specific areas indicated by arrows (HE, 400 X); panel 4. Lung of infected rhesus macaque with foci of mildly thickened alveolar walls lined by cuboidal epithelial cells with hyperchromatic nuclei representing type II pneumocyte hyperplasia (arrows) and few aggregates of macrophages in the alveolar lumina (asterisks) (HE, 400 X). (B) panel 1. Cervical lymph node of cynomolgus macaque J16017 with marked follicular lymphoid hyperplasia. Numerous lymphoid follicles in the cortex (square), paracortex (arrowhead) and medulla (asterisk) are indicated (HE, 50 X); panel 2. Mesenteric lymph node of rhesus macaque R14002 with marked paracortical lymphoid hyperplasia and medullar sinusoidal histiocytosis. Cortex (square), paracortex (arrowhead), and medulla (asterisk) are indicated (HE, 50 X). (C) panel 1. PET-CT image of the skull of J16017 with prominent metabolic activity in the left ocular region (arrow); panel 2. Conjunctiva of the same individual. Numerous perivascular and multifocal lymphocytes infiltrating the stratified squamous epithelium and the substantia propria. Goblet cell (arrow), non-keratinized stratified squamous epithelium (arrowhead), substantia propria (asterisk) are indicated (HE, 200X)

Rhesus macaque R14002 showed evidence of SARS-CoV-2 infection in the mesenteric lymph nodes by positive PCR and histologically exhibited marked lymphoid hyperplasia - paracortical type (Fig 8B panel 2).

## Discussion

In humans, COVID-19 displays a broad spectrum of disease symptoms [3, 59]. To investigate factors leading to the different COVID-19 manifestations animal models are essential as the moment and site of infection can be controlled and the subsequent response can be monitored in detail. Due to their similarity to humans, NHPs play a pivotal role in this type of preclinical research. To best appreciate the potential of the various macaque species as SARS-CoV-2 infection models, a thorough characterization of the entire course of infection is needed. The comparative study reported here contributes to this knowledge and further validates the different macaque models that are currently in use for COVID-19 research.

The study specifically focused on the post-acute phase of the SARS-CoV-2 infection in rhesus and cynomolgus macaques, defined as the 3-4 weeks period following the disappearance of detectable virus from the nose and throat swab samples. The majority of published COVID-19 NHP studies concentrate on the first two weeks of infection when evidence was found for acute viral interstitial pneumonia and virus was cleared from the nose and throat samples and lung lavages [29, 33, 35, 46, 60]. As the PET-CTs performed in this study on days 8 and 10 pi revealed an increase in ^18^F-FDG-uptake in the lungs and various tracheobronchial lymph nodes, it was decided to extend the observation period with weekly PET-CTs for all macaques for several weeks (rhesus macaques 35 or 36 days pi, for cynomolgus macaques 38 or 42 days pi, depending on the PET-CT findings).

During the entire study, including the acute infection phase, only mild clinical symptoms were noticed. This sharply contrasts with findings of others that described SARS-CoV-2-induced clinical disease, with clear signs of fever directly following infection, respiratory symptoms, and changes in hematology and clinical chemistry parameters [46, 48]. Unbiased measurement of the body temperature and activity of each animal was performed continuously by using telemetry. This is an important asset as in both macaque species a small, but notable elevation in body temperature was recorded in the first two weeks, the period of active virus replication. Significant differences in animal activity indicated that SARS-CoV-2 infection also influenced the well-being of the animals without causing obvious clinical symptoms. No indications for renal involvement or coagulation disorders were found in the macaques, and C-reactive protein (CRP) levels, a marker for pneumonia in humans [49], were also not afflicted. The clinical findings deviate from those described by other researchers. This discrepancy may be due, for instance, to the virus strain used for the challenge, the methods used for virus inoculation, the origin, genetic background, and environmental conditions of the animals.

In COVID-19 patients, the acute respiratory syndrome sometimes coincides with hypercytokinemia, occasionally resulting in multi-organ failure [61]. Patients with severe COVID-19 had significantly elevated plasma levels of proinflammatory cytokines [62]. In this study, macaques did not show overt disease symptoms but levels of certain cytokines, like IL-6, IFN-*γ*, MIP-1α, and MIP-1β increased in the plasma of both macaque species, showing that the chemokine system is involved in SARS-CoV-2 infection. The cytokine profiles after SARS-CoV-2 were highly comparable between the macaque species, except for IP-10 and MCP-1, suggesting differential involvement of monocyte activation between the species. We observed an increase in IL-6 levels at the end of the study in some individuals. In humans, IL-- 6 is involved in many inflammatory diseases. In patients with COVID-19, systemic production of IL-6 is associated with disease progression [63]. Also, in our study, the elevation of IL-6 corresponded with lung pathology as indicated by the highest CT/pathology scores in these animals. In humans, higher IL-6 levels coincide with increased CRP. We did not observe this correlation in the macaques, underlining subtle differences between human and NHP pathology in response to SARS-CoV-2 infection.

Antibody responses were detectable in all animals after infection. IgG titers to the N and S proteins, as well as the receptor-binding domain (RBD) on S, started to develop around day 14 and apart from individual differences continued to rise during the follow-up period. The development of IgG accurately followed the course of virus infection, as it became first detectable within one week after the virus had become undetectable in sera by RT-PCR. This was in line with findings of others [28, 47, 64]. No IgM directed to the N protein was detected in 7 out of 8 animals. This result was confirmed by a second, in-house developed IgM-ELISA (not shown) and thus a technical flaw in the serological assay used was excluded. We cannot explain the lack of N-specific IgM response to infection. In contrast, were measured elevated antibody levels against S and the RBD, most prominent in the cynomolgus macaques. Overall, IgM titers developed simultaneously with IgG, a salient phenomenon also reported by Hartman *et al*. in SARS-CoV-2-infected African Green monkeys [64].

Imaging was used to visualize possible lung pathology. In humans, CT is also applied, but the type of lung lesions found are only partly specific to COVID-19 [3, 65]. In the current study, purpose-bred NHPs with a well-documented health status were used, and pre-infection control scans were made. In this well-controlled NHP study, CT imaging is likely a valuable tool to monitor the progression of COVID-19-related lung pathology. Based on the criteria set to determine clinical severity [66], the macaques featured moderate disease levels as all eight individuals showed levels of pneumonia during the infection. Like in other NHP studies [28], lesions were observed in the lungs of infected animals. Differences in the location of lesions between the different studies may be related to different circumstances e.g. the method of administration of the virus into the trachea. Similar to Finch *et al*. [28], we did not collect bronchoalveolar lavage (BAL) samples in order to avoid unwanted interference with CT imaging [64, 67-69]. Instead, we collected tracheal swabs for viral load analysis. The viral RNA loads, but also the temporal pattern of RNA detection in swabs samples were similar to those observed in BAL [32, 35, 40, 41].

PET-CT imaging of the respiratory tract started in the late acute phase of infection and was regularly performed until euthanasia. What became evident, was that after assumed resolution of infection, as witnessed by negative testing for viral RNA in nasal and tracheal swabs, clear signs of pneumonia were still present in the lungs, with both sustained and newly formed lesions. Besides, an increased ^18^F-FDG uptake was detectable in the lungs and tracheobronchial lymph nodes of all animals. This indicates that despite mild disease symptoms in the acute phase of infection, and resolved viremia, SARS-CoV-2-induced pathogenesis continued. In addition, the detection of subgenomic messenger RNA in the respiratory tract and lung lymph nodes of 50% of animals implies that replication of SARS-CoV-2 continued unnoticed, and one can only hypothesize that this can cause disease symptoms in the macaques at a later stage. This finding in a recognized NHP model for COVID-19 research is of particular interest because of the growing concern for long-COVID in humans [70]. More than a year after the start of the pandemic, it is evident that humans can suffer from COVID-19-related symptoms, weeks to months after seemingly resolving the infection [5-7].

COVID-19 was initially regarded as a respiratory disease, but patients that succumb to this disease can display a complex array of pathologies that cover a broad spectrum of symptoms. In the macaques, viral RNA was also detected in other tissues, like in the conjunctiva, salivary gland, cervical and mesenteric lymph nodes, heart, and kidney. In COVID-19 patients these organs can also be affected by SARS-CoV-2, causing extra-pulmonary disease symptoms [59, 71, 72]. Histopathological findings of lymphocytic infiltrates in the lamina propria of the conjunctiva, and lymphoid hyperplasia in a cervical lymph node (both in J16017), as well as perivascular lymphocytic infiltrates in heart tissue and lesions in the mesenteric lymph node (both R14002), correlate with detection of viral RNA, and highlight the similarities in disease course between NHP and humans.

PET-CT imaging, in combination with viral RNA and subgenomic messenger RNA detection, revealed new insights in the post-acute phase of SARS-CoV-2 infection in NHP. Our data indicate widespread tissue dissemination of SARS-CoV-2 and provide clear evidence of continuing virus replication in lungs and surrounding lymph nodes after alleged convalescence of infection. This finding is intriguing as it has been hypothesized that persistent infection contributes to long COVID-19 in humans [70]. One wonders whether the current worldwide COVID-19 vaccination program will not only eliminate the acute disease but will have a positive effect on minimizing the long-COVID-related pathologies as well.

## Materials and Methods

### Ethics and Biosafety Statement

All housing and animal care procedures took place at the Biomedical Primate Research Centre (BPRC) in Rijswijk, the Netherlands. The BPRC is accredited by the American Association for Accreditation of Laboratory Animal Care (AAALAC) International and is compliant with European directive 2010/63/EU as well as the “Standard for Humane Care and Use of Laboratory Animals by Foreign Institutions” provided by the Department of Health and Human Services of the US National Institutes of Health (NIH, identification number A5539-01). Upon positive advice by the independent ethics committee (DEC-BPRC), the competent national authorities (CCD, Central Committee for Animal Experiments) issued a project license (license AVD5020020209404). Approval to start was obtained after further assessment of the detailed study protocol by the institutional animal welfare body (AWB) (in Dutch: Instantie voor Dierenwelzijn, IvD). All animal handlings were performed within the Department of Animal Science (ASD) according to Dutch law. ASD is regularly inspected by the responsible national authority (Nederlandse Voedsel-en Warenautoriteit, NVWA), and the AWB.

### Animals

Four Indian-origin rhesus macaques and four cynomolgus macaques were used in this study (Table S4). All macaques were mature, outbred animals, purpose-bred, and housed at the BPRC. The animals were in good physical health with normal baseline biochemical and hematological values. All were pair-housed with a socially compatible cage-mate in cages of at least 4 m^3^ with bedding to allow foraging and were kept on a 12-hour light/dark cycle. The monkeys were offered a daily diet consisting of monkey food pellets (Ssniff, Soest, Germany) supplemented with vegetables and fruit. Enrichment was provided daily in the form of pieces of wood, mirrors, food puzzles, and a variety of other homemade or commercially available enrichment products. Drinking water was available *ad libitum* via an automatic watering system. Animal Care staff provided daily visual health checks before infection, and twice-daily after infection. The animals were monitored for appetite, general behavior, and stool consistency. All possible precautions were taken to ensure the welfare and to avoid any discomfort to the animals. All experimental interventions (intratracheal and intranasal infection, swab collections, blood samplings, and (PET-)CTs were performed under anesthesia.

### Virus

The animals were infected with SARS-CoV-2 strain BetaCoV/BavPat1/2020. This strain was isolated from a patient who traveled from China to Germany, and an aliquot of a Vero E6 cell culture was made available through the European Virus Archive-Global (EVAg). The viral stock for the infection study was propagated on Vero E6 cells. For this study, a fifth passage virus stock was prepared with a titer of 3.2×10^6^ TCID_50_ per ml. The integrity of the virus stock was confirmed by sequence analysis.

### Experimental infections and post-exposure study follow-up

Three weeks before the experimental infection, a Physiotel Digital device (DSI Implantable Telemetry, Data Sciences International, Harvard Bioscience, UK) was implanted in the abdominal cavity of each animal. This device allowed the continuous real-time measurement of the body temperature and the animals’ activity remotely using telemetry throughout the study.

On day 0, all animals were exposed to a dose of 1 x 10^5^ TCID_50_ of SARS-CoV-2, diluted in 5 ml phosphate-buffered saline (PBS). The virus was inoculated via a combination of the intratracheal route, just below the vocal cords, (4.5 ml) and intranasal route (0.25 ml in each nostril). Virus infection was monitored for 35 to 42 days, and during that period the animals were checked twice daily by the animal caretakers and scored for clinical symptoms according to a previously published though adapted scoring system [73] (Table S2). A numeric score of 35 or more per observation time point was predetermined to serve as an endpoint and justification for euthanasia. Every time an animal was sedated, the body weight was measured. Blood was collected using standard aseptic methods from the femoral vein at regular time points post-infection (pi). In parallel, tracheal, nasal, and anal swabs were collected using Copan FLOQSwabs (MLS, Menen, Belgium). Swabs were placed in 1 ml DMEM, supplemented with 0.5% bovine serum albumin (BSA), fungizone (2.5 μg/ml), penicillin (100 U/ml), and streptomycin (100 μg/ml) and directly transported to the BSL3 lab.

### Biochemistry and hematology

Clinical biochemistry was performed using a Vetscan VS2 Chemical analyzer (Zoetis Benelux, Capelle aan de IJssel, Netherlands) with the use of the Comprehensive Diagnostic profile. This profile allows testing for alanine aminotransferase, albumin, alkaline phosphatase, amylase, calcium, creatinine, globulin, glucose, phosphorus, potassium, sodium, total bilirubin, total protein, and blood urea nitrogen. Hematology was done using a Vetscan HM5 Hematology analyzer (Zoetis Benelux, Capelle aan de IJssel, Netherlands). C-reactive protein and D-dimer levels were measured using Cobas Integra 400 plus analyzer (Roche Diagnostics Nederland B.V.).

### Detection of viral RNA in swabs, blood and tissue

To determine the presence of SARS-CoV-2 RNA in post-mortem tissues, tissue samples were weighed and placed in gentleMACS M tubes (30 mg in 100 μl PBS) and dissociated using a gentleMACS Tissue Dissociator (protein01 program)(Miltenyi Biotec B.V., Leiden, Netherlands). Next, the homogenized tissue was centrifuged for 10 min at 820 x g, and 100 μl supernatant was used for RNA isolation.

Viral RNA was isolated from plasma, swab sample supernatants, and cleared tissue homogenates using a QIAamp Viral RNA Mini kit (Qiagen Benelux BV, Venlo, Netherlands) following the manufacturer’s instructions. Viral RNA was reverse-transcribed to cDNA using a Transcriptor First Strand cDNA Synthesis kit (Roche Diagnostics BV, Almere, Netherlands). Viral genomic RNA was quantified by real-time quantitative RT-PCR specific for the RdRp gene of SARS-CoV-2, as described by Corman *et al*. [74]. The lower detection limit of the qRT-PCR was 3.6 viral RNA copies per reaction. Viral subgenomic messenger RNA (sgmRNA) was detected and quantified as previously described by Wölfel *et al* [75]. For both assays, RNA standard curves were generated by *in vitro* transcription of the target regions.

### Imaging

Positron Emission Tomography (PET)-computed tomography (CT) and CT data were acquired on multiple time points post-infection using a MultiScan Large Field of View Extreme Resolution Research Imager (LFER) 150 PET-CT (Mediso Medical Imaging Systems Ltd., Budapest, Hungary) as described before [76]. Animals fasted overnight and were sedated with ketamine (10 mg/kg ketamine hydrochloride (Alfasan Nederland BV, Woerden, Netherlands) combined with medetomidine hydrochloride (0.05 mg/kg (Sedastart; AST Farma B.V., Oudewater, Netherlands)) to induce sedation and muscle relaxation, both applied intramuscularly (IM). The animals were positioned head-first supine (HFS) with the arms up. After the scan, upon return to their home cage, atipamezole hydrochloride (Sedastop, ASTFarma B.V., Oudewater, Netherlands, 5 mg/ml, 0.25 mg/kg) was administrated IM to antagonize medetomidine.

#### PET-CT

The PET-CT images were acquired under mechanical ventilation in combination with a forced breathing pattern. For anesthetic maintenance, a minimum alveolar concentration of isoflurane (iso-MAC) of around 0,80%-1.00% was used. The body temperature of the animal was maintained by using the Bair Hugger (3M^TM^, St Paul, MN, USA) supplied with 43°C airflow. Typically, around 100 MBq of ^18^F-FDG was applied intravenously (GE Healthcare, Leiderdorp, Netherlands). A 15-minute static PET was acquired of the lungs 45 minutes after injection. Afterward, the emission data was iteratively reconstructed (OSEM3D, 8 iterations, and 9 subsets) into a single frame PET image normalized and corrected for attenuation, scatter, and random coincidences using the reference CT and corrected for radioactive decay. The analysis was performed in VivoQuant 4.5 (Invicro, Boston, USA). Lung lesions were discriminated based on lung density as defined by Hounsfield Units (HU) on CT. Lymph node uptake was quantified with the use of a lower threshold standard uptake value (SUV) of 2.0 to discriminate activated lymph nodes from the surrounding tissue. Two anatomical and two molecular output parameters were generated; volume, anatomical density in HU, average standardized uptake value (SUVmean), and a maximum SUV corrected for scatter and random coincidences (SUVpeak). In this way, a full overview of the results could be obtained with minimal inter and intra-observer variation [76].

#### Gated-CT

In order to mitigate motion artefacts and to allow a non-invasive CT, retrospective gating was applied during CT. The respiratory amplitude was detected with a gating pad placed next to the umbilicus. For the final reconstruction, the inspiration phases were exclusively used and manually selected. A semi-quantitative scoring system for chest CT evaluation was used to estimate SARS-CoV-2-induced lung disease [28, 55, 67]. Quantification of the CTs was performed independently, by two experienced imaging scientists based on the lobes; the middle and accessory lobe of the right lung were combined. The degree of involvement in each zone was scored as: 0 for no involvement, 1 for <5%, 2 for 5-24%, 3 for 25-49%, 4 for 50-74% and 5 for >74% involvement. An additional increase or decrease of 0.5 was used to indicate alterations in CT density of the lesions. By using this scoring system, a maximum score of 30 could be reached for the combined lobes per time point.

### Cytokine and chemokine analysis

Cytokine and chemokine concentrations in sera of infected macaques, including IL-1β, IL-6, CCL11 (Eotaxin), CXCL10 (IP-10), CXCL11 (I-TAC), CCL2 (MCP-1), CXCL9 (MIG), CCL3 (MIP-1*α*), CCL4 (MIP-1β), CCL5 (RANTES), CXCL8 (IL-8), TNF-*α*, and IFN-*γ*, were determined using LEGENDplex™ NHP Chemokine/Cytokine Panel (13-plex) (Biolegend, San Diego, CA, USA) essentially according to manufacturer’s instruction. Samples were measured on an LSRII FACS machine (BD Biosciences, Vianen, Netherlands) and analyzed by using company software.

### Serology

Antibodies to SARS-CoV-2 in sera were analyzed using different enzyme-linked immunosorbent assays. A double recognition enzyme-linked immunosorbent assay (DR-ELISA) that detects total immunoglobulins targeted to the SARS-CoV-2 nucleoprotein (N) protein was used, as described by Hoste *et al.* [77] (INgezim COVID19 DR; Eurofins-INGENASA, Madrid, Spain). In addition, an ELISA that detects immunoglobulins elicited to the full-length spike (S) protein, and an assay that specifically detects antibodies directed to the receptor-binding domain (RBD) of the S protein were applied in this study.

For the analysis of anti-S protein antibody responses, Greiner half-area ELISA plates were coated overnight with 1 µg/ml of the monomeric full-length Spike or Spike RBD proteins (both antigens expressed in insect cells; AdapVac, Copenhagen, Denmark) in PBS. Plates were washed and blocked for one hour with PBS Tween-20 (0.05%) + BSA (3% m/v). Serial dilutions of sera were made in PBS Tween-20 (0.05%) + BSA (1% m/v) (PBS-TB) and added to the plate in duplicate. As a standard reference, a monoclonal antibody specific for the SARS-CoV-2 RBD was used, diluted in PBS-TB to an initial IgG concentration of 500 ng/ml, and diluted in the ELISA plate in a two-fold series over seven wells (Range 500 to 0.007813 ng/ml). Standards were run in duplicate. The reference standard and sera were incubated for one hour at 37°C, and subsequently washed with PBS Tween-20 (0.05%). Goat-anti-human (H+L) IgG-HRP (Thermo Fisher Scientific, Waltham, MA, USA) was added to the plate and incubated for one hour at 37°C after which the plates are washed and TMB substrate was added. The reaction was stopped after 15 min by the addition of 0.2 M H_2_SO_4_. Plates were then read at 450 nm on a SpectraMax M5 plate reader (Molecular Devices, LLC., San Jose, CA, USA), and OD values were saved as Excel files. A four-parameter curve was fit to the standard, and IgG concentrations in the samples were calculated from the OD values using the parameters as estimated from the four-parameter fit. Only OD values in the linear part (i.e. parallel to the standard curve) of the dilution curve were used for the samples. Within sample coefficient of variation (CV) for the included OD values usually was below 25%.

To detect IgM, the same protocol was followed, but the secondary antibody used was an anti-human IgM Peroxidase-labelled antibody produced in goats (SouthernBiotech, Birmingham, USA).

### Necropsy and histological analysis

After euthanasia necropsies were performed according to a standard protocol. The samples were fixed by immersion in 10% neutral-buffered formalin, routinely processed into paraffin blocks, cut into 4 microns tissue sections, stained with hematoxylin and eosin (H&E), and examined microscopically. In parallel, samples from the same organs were collected for virus detection by real-time PCR and for IHC analyses. Additional special staining (Gram-staining) for the presence of bacteria was done on all seven pulmonary lobes of each animal and was confirmed negative.

## Acknowledgements

We would like to thank the Animal Science Department, especially the animal caretakers and veterinarians for excellent care of the animals. Francisca van Hassel for her assistance in figure design. We also thank Cristina Aira (Eurofins Ingenasa, Madrid) for technical assistance.

## Supporting information

**Fig S1.**
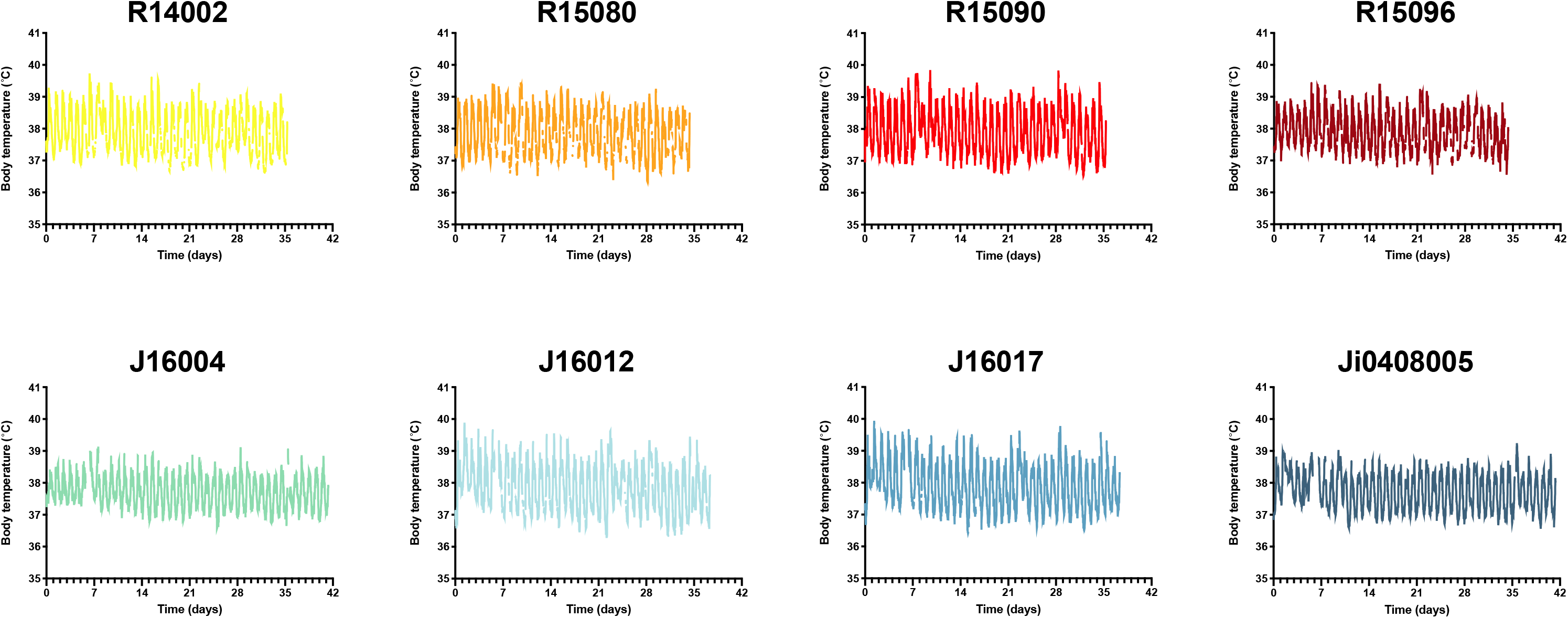
Body temperatures of all animals. Body temperatures were measured using a digital telemetric device during the entire study. The gaps in the graphs are due to data loss during measurement.

**Fig S2.**
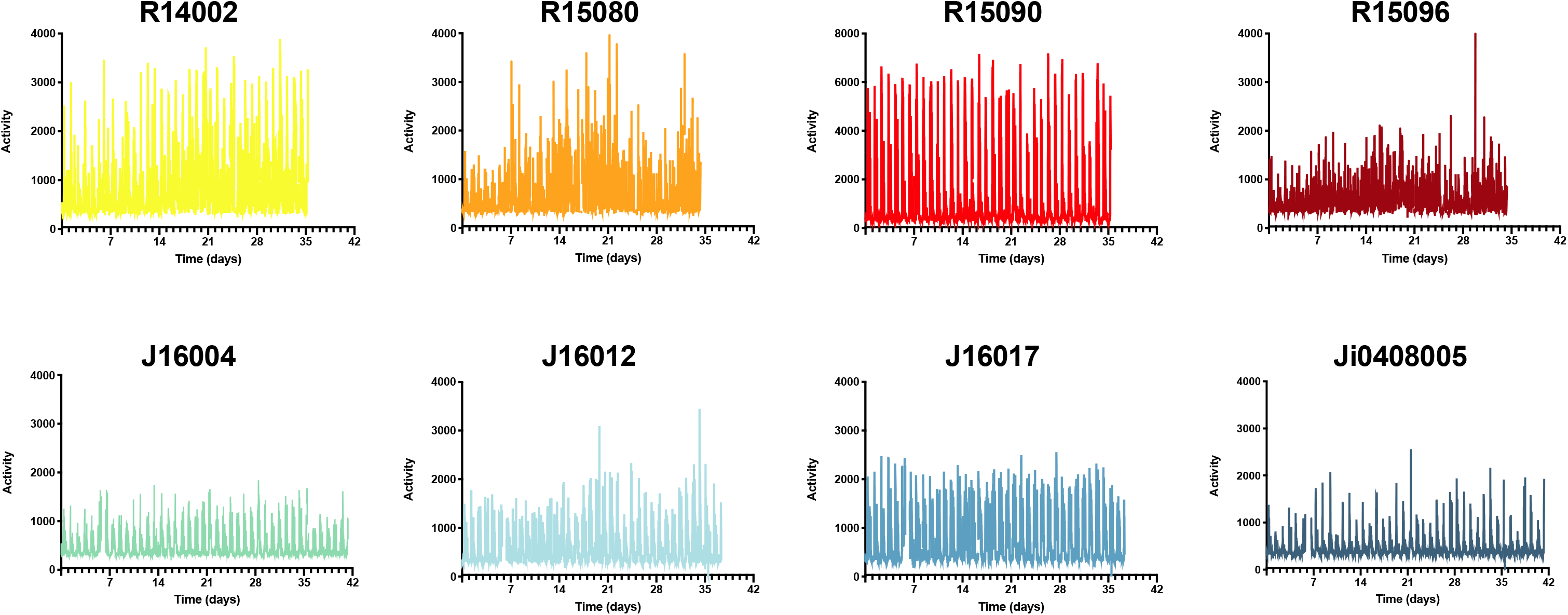
Activity of all animals. (A) The activity of each animal was measured using a digital telemetric device during the entire study. Gaps in the graphs are due to data loss during measurement. (B) Cumulative activity scores of the first two weeks compared to the third week of the study, calculated as total area under the curve for the given period.

**Fig S3.**
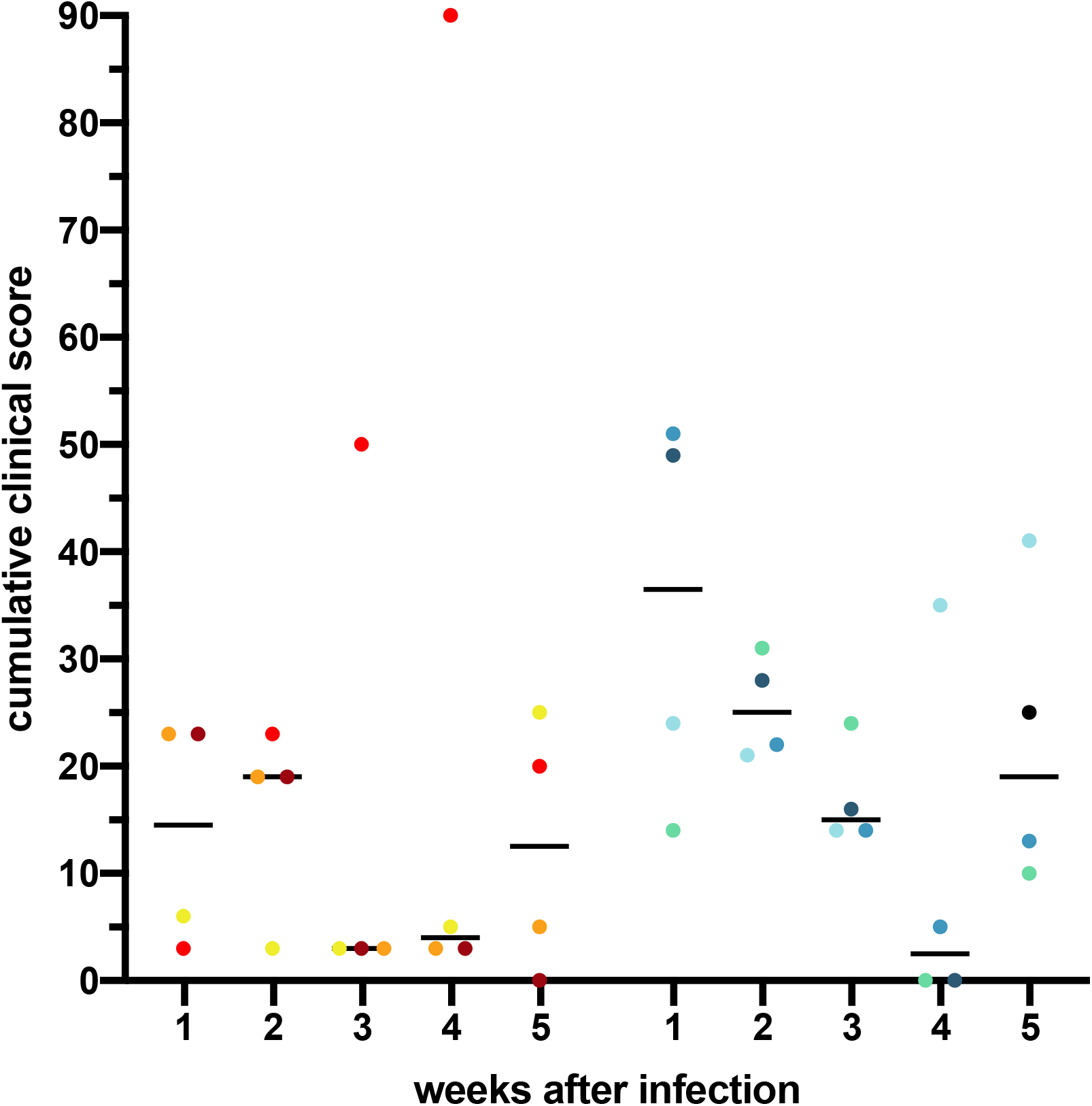
Cumulative clinical scores. Scores were calculated per week and per individual animal. Horizontal bars represent medians.

**Fig S4.**
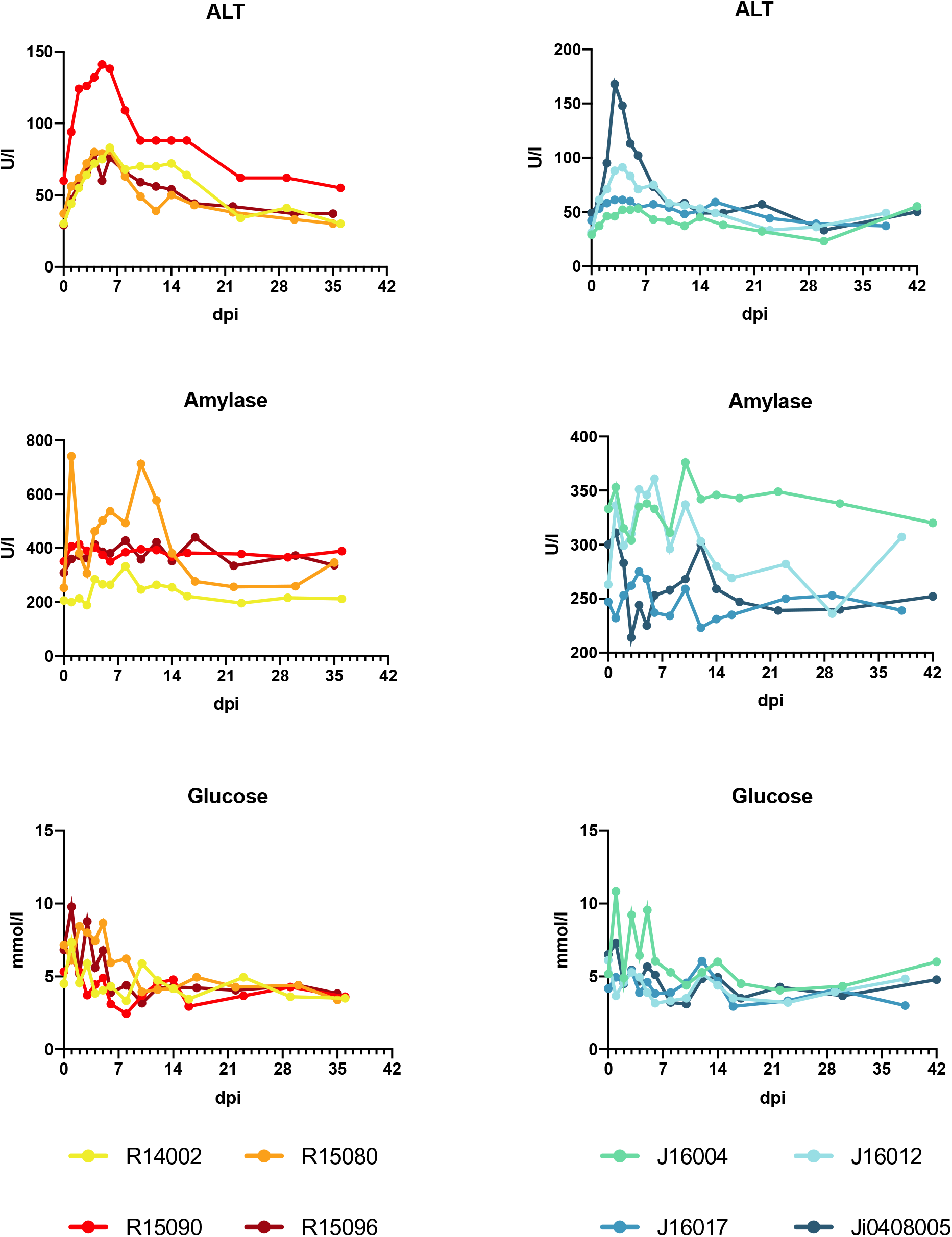
Clinical biochemistry. ALT (alanine aminotransferase), amylase and glucose levels were measured in serum samples of rhesus and cynomolgus macaques

**Fig S5.**
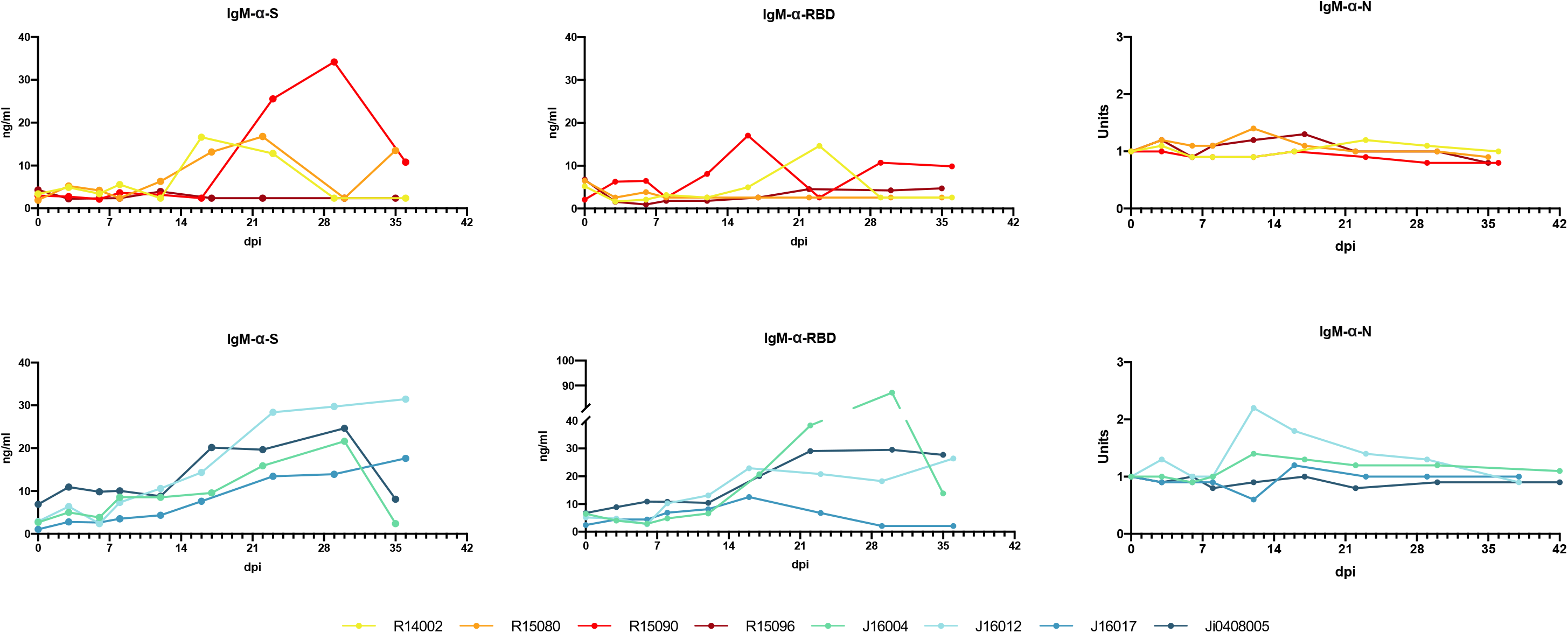
Development of SARS-CoV-2 IgM response in rhesus and cynomolgus macaques. The IgM response in serum was determined using an anti-S IgM ELISA, a serological test to detect IgM directed to the RBD, and an anti-N IgM ELISA (left to right).

**Fig S6.**
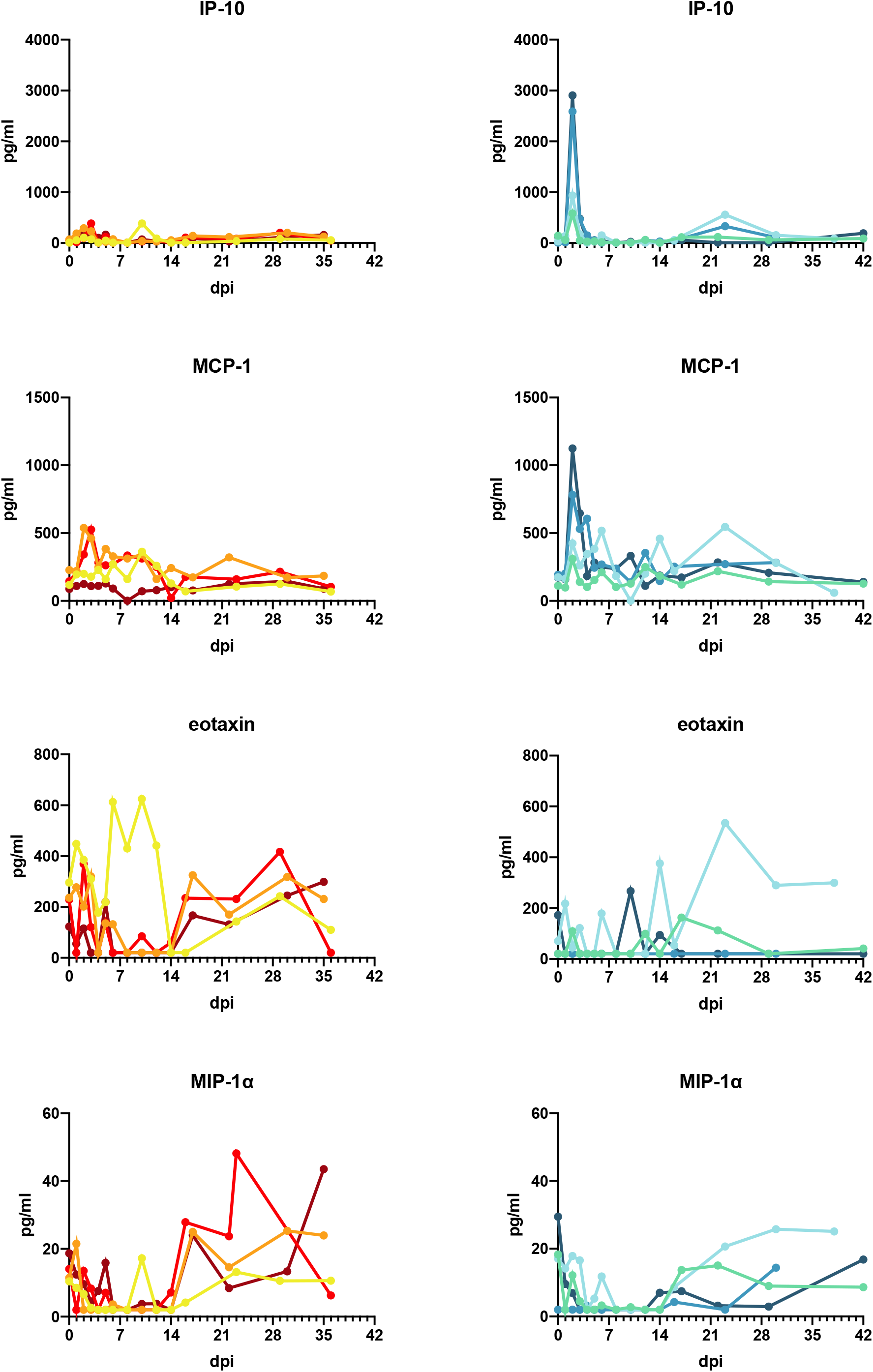
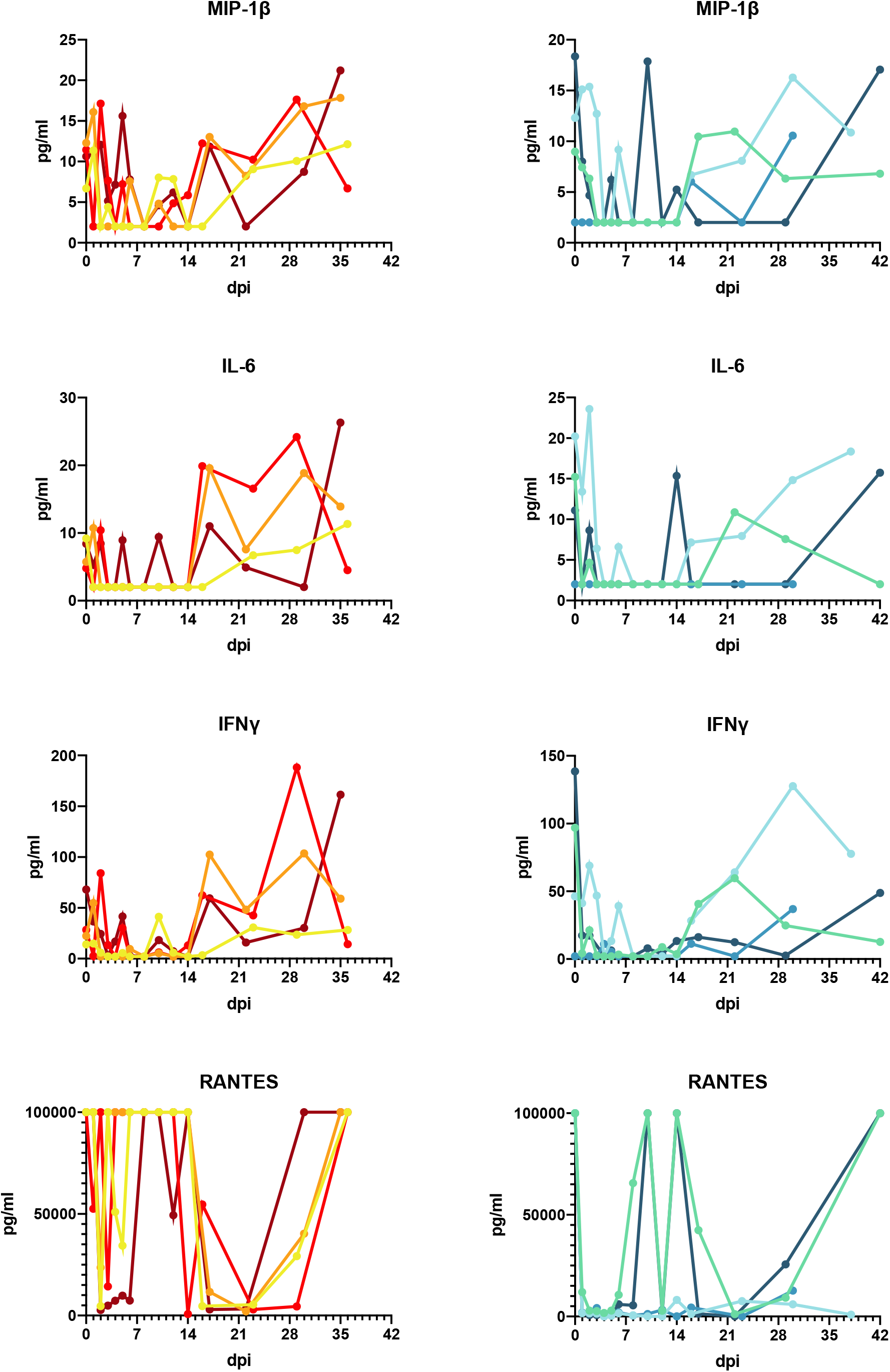
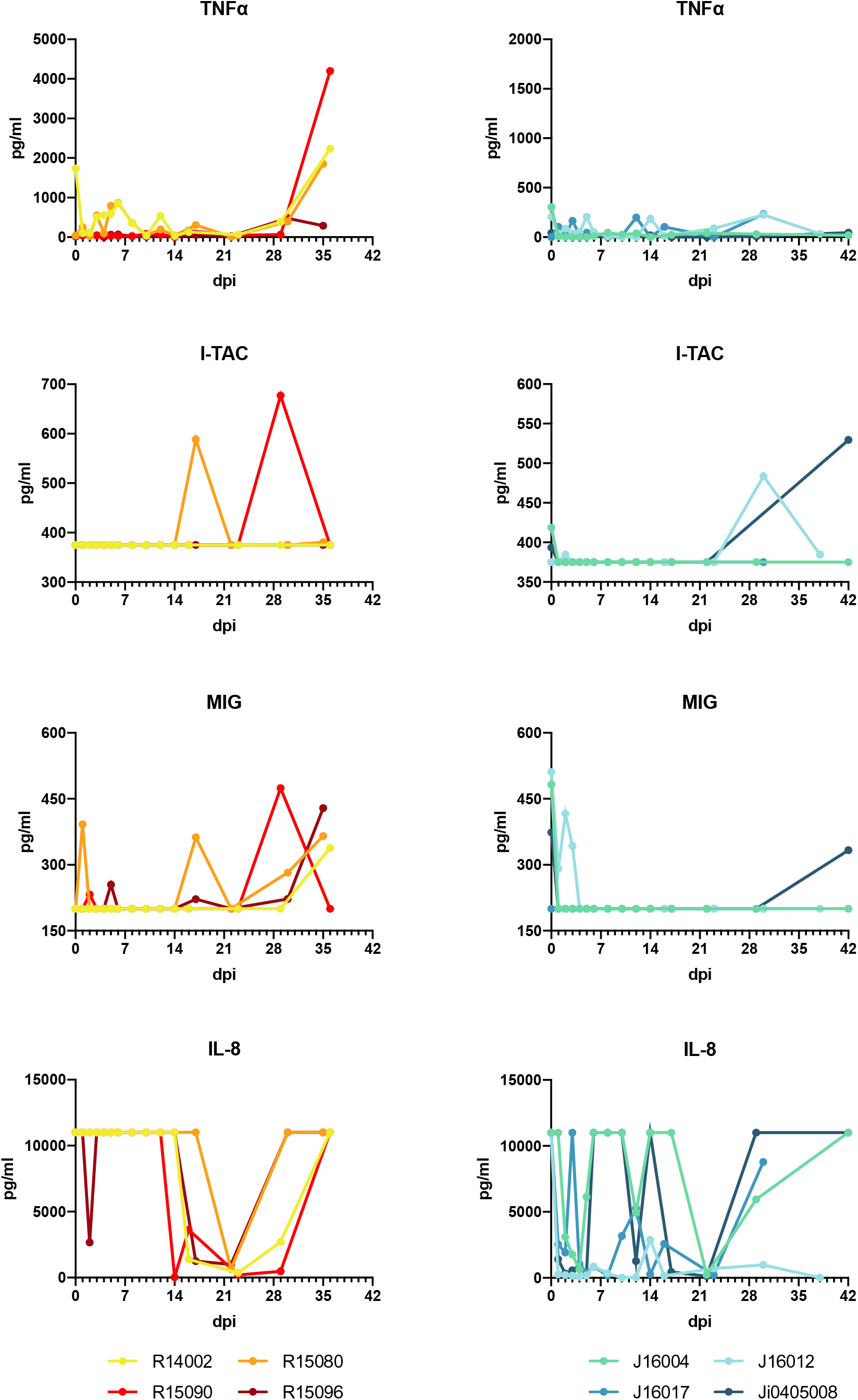
Cytokine and chemokine levels in SARS-CoV-2-infected macaques. Levels were determined using LEGENDplex™ NHP Chemokine/Cytokine Panel (13-plex). Samples were measured on a LSRII FACS machine and analyzed by using company software.

**Fig S7.**
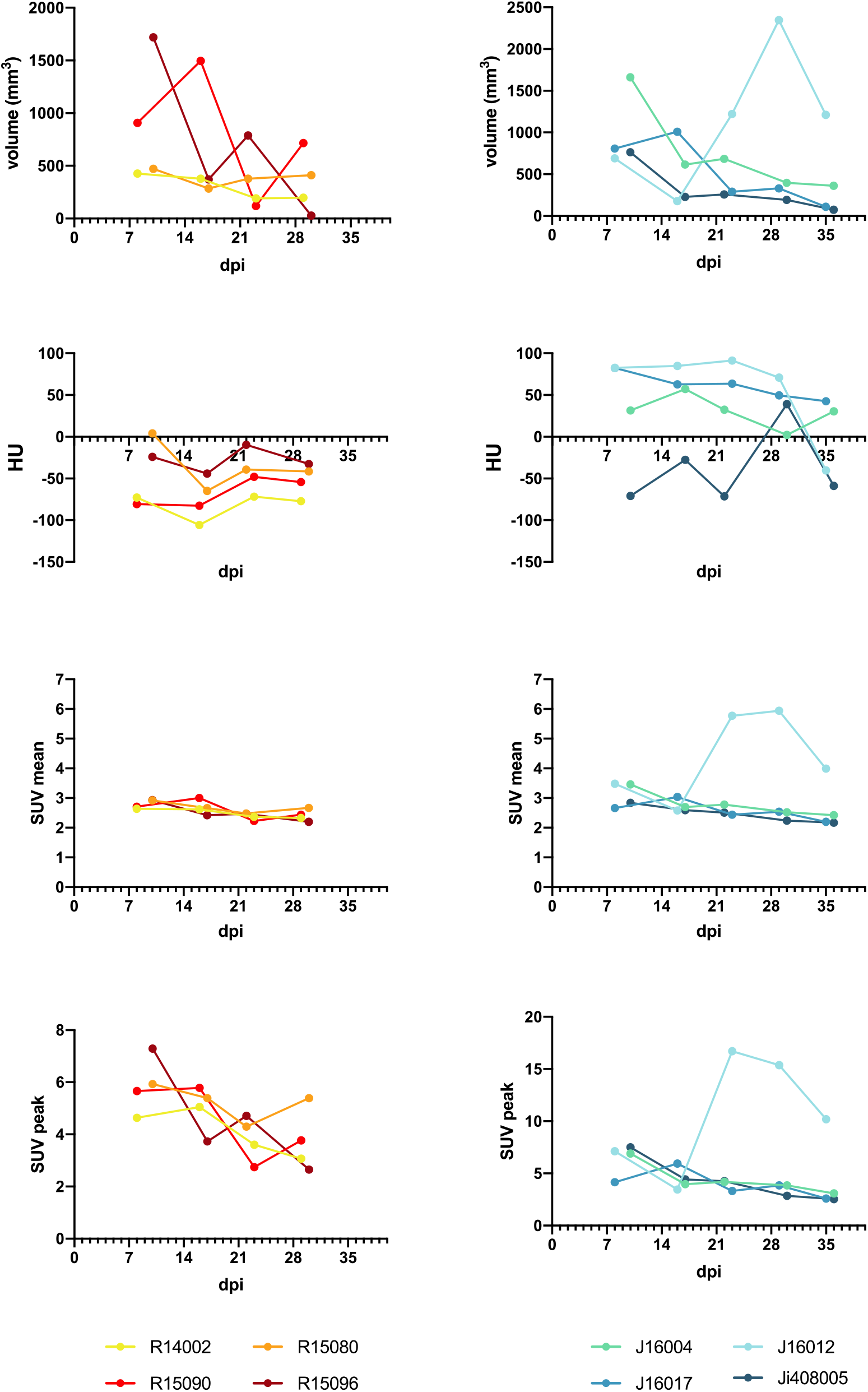
Quantification of ^18^F-FDG uptake by tracheobronchial lymph nodes. The tracheobronchial lymph nodes were quantified on PET-CT with two anatomical (upper rows) and two metabolic parameters (bottom rows). The anatomical parameters are; volume in mm^3^ and density in Hounsfield Units (HUs). The metabolic parameters are the average metabolic uptake represented by the mean standard uptake value (SUV) and the maximum uptake in a region of interest (ROI), corrected for random and scattered coincidences represented by the SUVpeak.

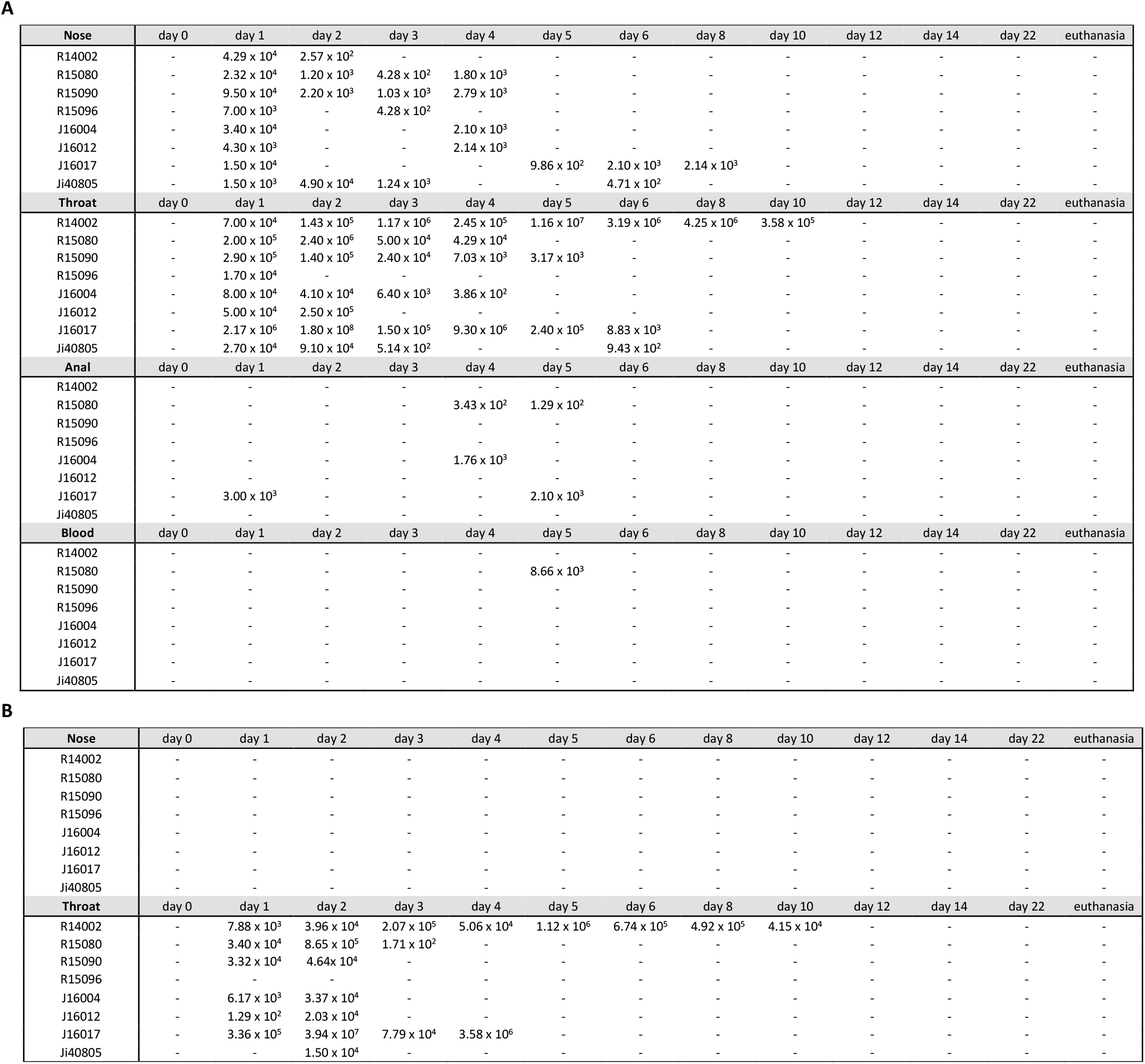
**Table S1A.** Viral loads in swab and blood samples (RNA genome equivalents/ml) **Table S1B.** Subgenomic messenger RNA in tracheal and nasal swab samples (sgmRNA copies/ml)

**Table S2.**
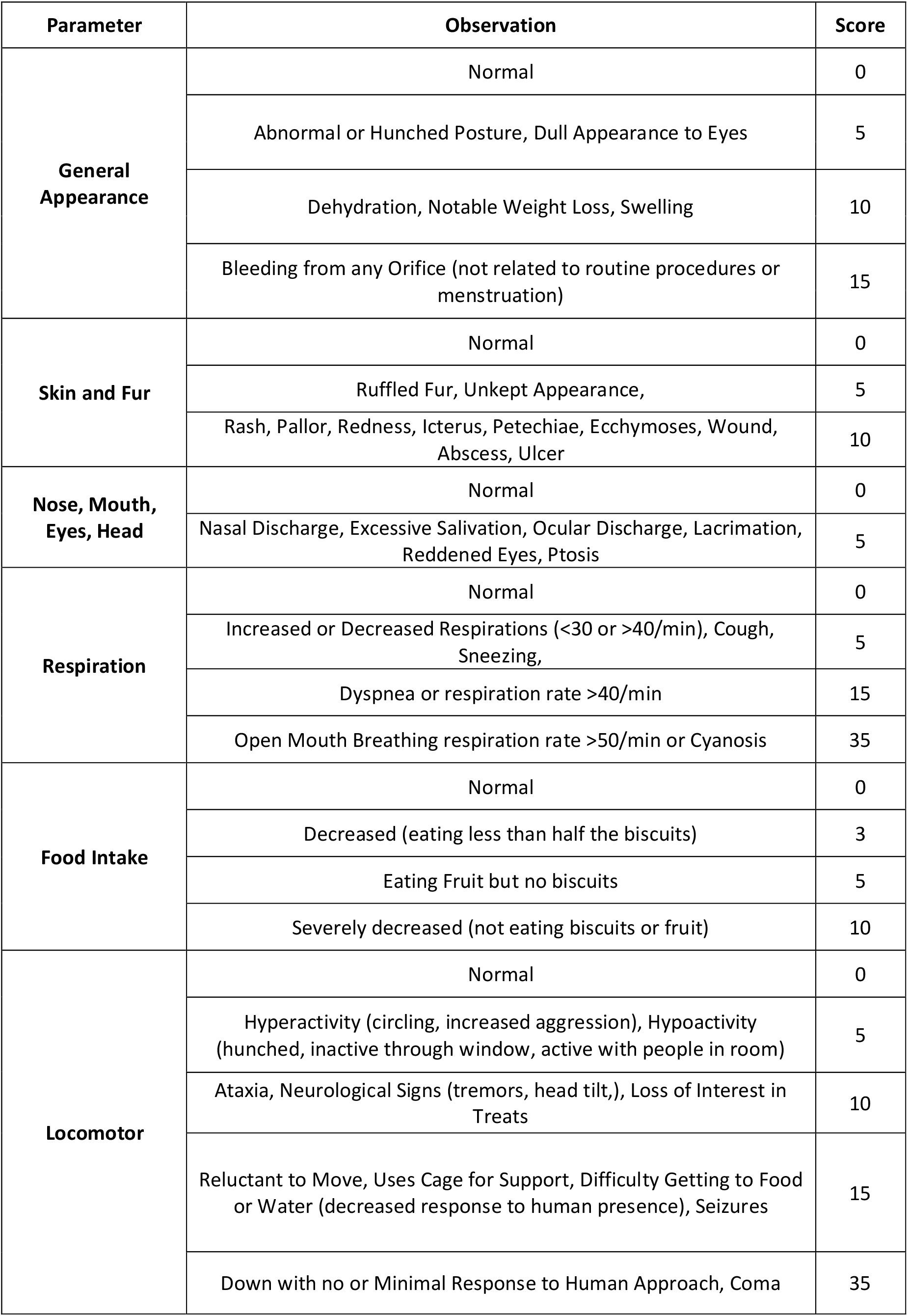
Clinical symptoms scoring list. Adapted to Brining *et al.* 2010, Comparative Medicine, Vol 6, No 5, pp 389-396.

| Parameter | Observation | Score |
| --- | --- | --- |
| **General Appearance** | Normal | 0 |
|  | Abnormal or Hunched Posture, Dull Appearance to Eyes | 5 |
|  | Dehydration, Notable Weight Loss, Swelling | 10 |
|  | Bleeding from any Orifice (not related to routine procedures or menstruation) | 15 |
| **Skin and Fur** | Normal | 0 |
|  | Ruffled Fur, Unkept Appearance, | 5 |
|  | Rash, Pallor, Redness, Icterus, Petechiae, Ecchymoses, Wound, Abscess, Ulcer | 10 |
| **Nose, Mouth, Eyes, Head** | Normal | 0 |
|  | Nasal Discharge, Excessive Salivation, Ocular Discharge, Lacrimation, Reddened Eyes, Ptosis | 5 |
| **Respiration** | Normal | 0 |
|  | Increased or Decreased Respirations (<30 or >40/min), Cough, Sneezing, | 5 |
|  | Dyspnea or respiration rate >40/min | 15 |
|  | Open Mouth Breathing respiration rate >50/min or Cyanosis | 35 |
| **Food Intake** | Normal | 0 |
|  | Decreased (eating less than half the biscuits) | 3 |
|  | Eating Fruit but no biscuits | 5 |
|  | Severely decreased (not eating biscuits or fruit) | 10 |
| **Locomotor** | Normal | 0 |
|  | Hyperactivity (circling, increased aggression), Hypoactivity (hunched, inactive through window, active with people in room) | 5 |
|  | Ataxia, Neurological Signs (tremors, head tilt,), Loss of Interest in Treats | 10 |
|  | Reluctant to Move, Uses Cage for Support, Difficulty Getting to Food or Water (decreased response to human presence), Seizures | 15 |
|  | Down with no or Minimal Response to Human Approach, Coma | 35 |

**Table S3.** Viral RNA detection in organs of SARS-CoV-2 infected animals. Amount of viral RNA is given as genome equivalent per gram of tissue. Only tissue samples that were PCR-positive in at least one of the animals are shown in the table. *Number of lymph nodes sampled from the respiratory tract varied between individual animals. Nd; not done.

|  | R14002 | R15080 | R15090 | R15096 | J16012 | J16017 | J408005 | J16004 |
| --- | --- | --- | --- | --- | --- | --- | --- | --- |
| Skin lesion | - | - | - | - | - | 1 x 10<sup>4</sup> | - | - |
| Conjunctiva | - | - | - | - | - | 3.88 x 10<sup>4</sup> | - | - |
| Tongue mucosa | - | - | - | - | - | - | - | - |
| Nasal mucosa | - | - | - | - | - | - | - | - |
| Pharyngeal mucosa | - | - | - | - | - | 7.7 x 10<sup>4</sup> | 3.6 x 10<sup>3</sup> | - |
| Laryngeal mucosa | - | - | - | - | - | 1.23 x 10<sup>5</sup> | - | - |
| Deep cervical lymph node (LN) | - | - | - | - | - | 3.4 x 10<sup>3</sup> | - | - |
| tonsil | - | - | - | - | - | - | - | - |
| Mesenteric LN | 3.08 x 10<sup>4</sup> | - | - | - | - | 4.8 x 10<sup>3</sup> | - | - |
| heart, right atrium | - | - | - | - | - | 4.76 x 10<sup>4</sup> | - | - |
| heart, left atrium | 8.82 x 10<sup>5</sup> | - | - | - | - | - | - | - |
| heart, right ventricle | 1.62 x 10<sup>5</sup> | - | - | - | - | 1.15 x 10<sup>5</sup> | - | - |
| heart, left ventricle | - | - | - | - | - | 3.52 x 10<sup>4</sup> | - | - |
| liver | - | - | - | - | - | 5.93 x 10<sup>2</sup> | - | - |
| spleen | - | - | - | - | - | 2.13 x 10<sup>3</sup> | - | - |
| kidney | - | - | - | - | - | 1.08 x 10<sup>4</sup> | - | - |
| jejunum | - | - | - | - | - | - | - | - |
| ileum | - | - | - | - | - | - | - | - |
| colon | - | - | - | - | - | - | - | - |
| Salivary gland | - | - | - | - | - | 1.38 x 10<sup>4</sup> | - | - |
| carina | 8.8 x 10<sup>3</sup> | - | - | - | 4.4 10<sup>3</sup> | - | - | - |
| lung, upper right lobe | - | - | - | - | 2.2 x 10<sup>3</sup> | - | 5.2 x 10<sup>3</sup> | - |
| lung, accessory lobe | - | - | - | - | 3.08 x 10<sup>3</sup> | - | - | - |
| lung, middle right lobe | 2.74 x 10<sup>4</sup> | - | - | - | - | - | - | - |
| lung, lower right lobe | 4.28 x 10<sup>4</sup> | - | - | - | - | - | - | - |
| lung, upper left lobe | 2.2 x 10<sup>4</sup> | - | - | - | - | - | - | - |
| lung, middle left lobe | 1.9 x 10<sup>4</sup> | - | - | - | - | - | - | - |
| lung, lower left lobe | 6.59 x 10<sup>4</sup> | - | - | - | - | - | - | - |
| Calf vein | - | - | - | - | - | 1.5 x 10<sup>4</sup> | - | - |
| Bronchus left | - | - | - | - | - | 1.5 x 10<sup>4</sup> | - | - |
| Bronchus right | - | - | - | - | - | 3.2 x 10<sup>3</sup> | - | - |
| trachea | - | - | - | - | - | 7.43 x 10<sup>4</sup> | - | - |
| Left paratracheal LN* | 6.43 x 10<sup>5</sup> | - | - | - | - | 2 x 10<sup>5</sup> | nd | - |
|  | nd | nd | nd | nd | nd | 8.24 x 10<sup>5</sup> | nd | nd |
| Left hilar (bronchial) LN* | 9.82 x 10<sup>5</sup> | - | - | - | 7.47 x 10<sup>3</sup> | 2.58 x 10<sup>4</sup> | - | - |
|  | 4.72 x 10<sup>3</sup> | - | - | - | - | 6.02 x 10<sup>4</sup> | - | - |
|  | 5.67 x 10<sup>4</sup> | - | nd | - | - | nd | - | nd |
|  | - | nd | nd | - | - | nd | - | nd |
|  | nd | nd | nd | - | nd | nd | - | nd |
| Subcarinal LN* | - | - | - | - | 2.58 x 10<sup>4</sup> | 1.14 x 10<sup>4</sup> | - | - |
|  | nd | - | - | nd | nd | nd | nd | nd |
|  | nd | nd | - | nd | nd | nd | nd | nd |
|  | nd | nd | - | nd | nd | nd | nd | nd |
| Right hilar (bronchial) LN* | 9.61 x 10<sup>4</sup> | - | - | - | 5.65 x 10<sup>4</sup> | 8.52 x 10<sup>4</sup> | 4.11 x 10<sup>4</sup> | - |
|  | - | - | nd | - | - | nd | - | - |
|  | - | - | nd | nd | - | nd | - | nd |
|  | nd | nd | nd | nd | nd | nd | 8.1 x 10<sup>2</sup> | nd |
| Right paratracheal LN* | nd | nd | - | nd | 5.39 x 10<sup>4</sup> | 9.81 x 10<sup>4</sup> | nd | nd |
|  | nd | nd | - | nd | nd | 6.47 x 10<sup>3</sup> | nd | nd |

**Table S4.** List of animals used in the study. Rh: rhesus macaque, Cy: cynomolgus macaque. ^#^at the start of the study

| Species | monkey ID | Age | Body weight<sup>#</sup> | Sex |
| --- | --- | --- | --- | --- |
| R | R14002 | 6y | 8.2 kg | male |
| R | R15080 | 5y | 7.9 kg | male |
| R | R15090 | 5y | 7.8 kg | male |
| R | R15096 | 5y | 8.7 kg | male |
| C | J16004 | 4y | 5.7 kg | male |
| C | J16012 | 4y | 3.3 kg | male |
| C | J16017 | 4y | 4.9 kg | male |
| C | Ji0408005 | 16y | 9.7 kg | male |

